# Small Molecule Protein Assembly Modulators with Pan-Cancer Therapeutic Efficacy

**DOI:** 10.1101/2022.09.28.509937

**Authors:** Anuradha F. Lingappa, Olayemi Akintunde, Connie Ewald, Markus Froehlich, Niloufar Ziari, Maya Michon, Shao Feng Yu, Suguna Mallesh, Jim Lin, Anatoliy Kitaygorodskyy, Dennis Solas, Jonathan C. Reed, Jaisri R. Lingappa, Andreas Mueller-Schiffmann, Carsten Korth, Dharma Prasad, Aysegul Nalca, Emily Aston, Brad Fabbri, Sanjeev Anand, Thomas W. Campi, Emma Petrouski, Debendranath Dey, David W. Andrews, Vishwanath R. Lingappa

## Abstract

Two structurally-unrelated small molecule chemotypes, represented by compounds PAV-617 and PAV-951 with antiviral activity in cell culture against monkeypox virus (MPXV) and human immunodeficiency virus (HIV) respectively, were studied for anti-cancer efficacy. Each exhibited apparent pan-cancer cytotoxicity, reasonable pharmacokinetics, and non-toxicity in mice at active concentrations. Anti-tumor properties of both chemotypes, were validated in mouse xenografts against A549 human lung cancer and, for one of the chemotypes, against HT-29 colorectal cancer. The targets of these compounds are unconventional: each binds to a different transient, energy-dependent multi-protein complex containing the protein TRIM28/KAP1, an allosteric modulator known to regulate mechanisms underlying viral and nonviral disease states including cancer. Treatment with these compounds alters the target multi-protein complexes in a manner consistent with allosteric modulation as their mechanism of action. These compounds appear to remove a block, crucial for cancer survival and progression, on the homeostatic linkage of uncontrolled cellular proliferation to apoptosis. These compounds provide starting points for development of next-generation non-toxic, pan-cancer therapeutics.

## Background

The similarity of the interactions of viral infection and cancer with the healthy host has often been noted (1). Both represent pathological processes of extremely diverse origin, that overcome the complex feedback controls of homeostasis, to the detriment of the host (2). Both exploit natural selection as a powerful weapon to overcome host defenses. Viruses have done so over deep evolutionary time through co-evolution with their hosts and through the emergency of resistance mutations in response to the selective pressure of treatments (3,4). Cancers regularly use the latter mechanism, resulting in clonal mutants that drive cancer progression (5).

At least seven different viruses--Epstein-Barr virus (EBV), hepatitis B virus (HBV), hepatitis C virus (HCV), human T-lymphotropic virus 1 (HTLV-1), human papillomavirus (HPV), Kaposi sarcoma-associated herpesvirus (KHSV or HHV-8), and Merkel cell polyomavirus (MCPyV)—are known to be directly oncogenic through their alteration of the cellular environment and/or impairment of the host’s innate immune system defenses (6,7). It has therefore been proposed that viruses could play a key role in the discovery of new cancer treatments by identifying cellular targets that drive tumorigenesis (8).

The incompleteness in our current understanding of the dynamics of host homeostasis and its myriad of feedback controls has been a disadvantage for efforts to design novel therapeutic countermeasures against both viruses and cancer. However, a recent unconventional approach to antiviral drug discovery has identified small molecules targeting host allosteric sites essential for homeostasis, that are repurposed by viral infection (9). The antiviral compounds identified by this approach appear to restore key features essential to host homeostasis (9). We wished to determine whether those compounds might have therapeutic applicability against cancer, given the analogy in how both viruses and cancers drive departure from homeostasis. The results to be presented here suggest this is the case and shed light on molecular pathways relevant for both viral and neoplastic disease, providing a strategy for development of novel cancer cell therapeutics.

Utilizing a cell-free protein synthesis and assembly (CFPSA) system, compounds from a library of approximately 150,000 drug-like small molecules were screened for hits which blocked the assembly of viral capsids without inhibiting protein synthesis (9–12). Hit compounds identified have been termed “protein assembly modulators”. A collection of 300 structurally-diverse protein assembly modulators were identified from hits demonstrating activity against capsid assembly in one or more viral family (9). Antiviral protein assembly modulators from this collection have been validated against infectious virus in cell culture for multiple viral families including *Retroviridae*, *Rhabdoviridae*, *Poxviridae*, *Adenoviridae*, *Herpesviridae*, *Paramyxoviridae*, *Coronaviridae*, *Orthomyxoviridae*, and *Picornaviridae* (9,11–13). In two cases, *Coronaviridae* and *Paramyxoviridae*, cellular antiviral activity has been confirmed in animal disease models (9).

These antiviral protein assembly modulators appear to target host-viral protein-protein interactions via allosteric sites that control repurposing of host assembly machinery for viral capsid formation and which also allow disengagement of host innate immune defenses such as autophagy (9). In an earlier study, a class of protein assembly modulators was shown to change the composition of a transient, energy-dependent multi-protein complex whose components include p62/SQSTM1, a key regulator of autophagy (9). This multi-protein complex is co-opted upon viral infection and modified in composition. p62/SQSTM1 is lost from the complex, and the viral nucleoprotein added (9). Upon treatment with an antiviral protein assembly modulator, the target multi-protein complex is restored to its composition in uninfected cells, with loss of the viral nucleoprotein component and restoration of p62/SQSTM1 (9).

The discovery of dynamic multi-protein complexes whose composition changes with drug treatment offers a new means of parsing out post translational protein heterogeneity and its relevance for diseased states. The amount of a given protein that has been observed in the multi-protein complexes targeted by antiviral assembly modulators comprises only a very small fraction of the total amount of the component proteins present in a cell (9). The role played by the subset of the protein that is part of a particular multi-protein complex may reflect a “moonlighting” function, as observed for a growing number of cellular and viral proteins (14–16).

We hypothesized that if an overlap between viral and oncogenic pathways exists, some antiviral assembly modulators might be capable of disrupting a multi-protein complex associated with a hallmark of cancer (17,18). To test the hypothesis, we established a cancer-relevant counter screen and applied it to previously-identified antiviral protein assembly modulator compounds. In this paper, we will describe two protein assembly modulators which were originally characterized for their antiviral properties but are now shown to have potential as cancer therapeutics based on *in vitro* screening and *in vivo* validation studies. Just as the previously cited study discovered a pan-respiratory antiviral chemotype that appears to target a host-viral interface, so also we hypothesized that it should be possible to find a comparable anti-cancer chemotype that reverses changes selected by the cancer for allowing it to escape from feedback constraints that normally prevent uncontrolled proliferation.

## Results

### Uncontrolled cellular proliferation: a hallmark of cancer inhibited by protein assembly modulators

No hallmark of cancer is more fundamental than uncontrolled proliferation (18). Uncontrolled proliferation normally triggers cell death mechanisms, including apoptosis (19). Therefore, to survive a cancer must achieve a means of evading cell death long enough to complete cell division and reset the cell death timer. This, in turn, allows further proliferation, during which time additional mutations can occur and selection pressure will drive higher and higher grade malignancy and so on, ultimately resulting in metastasis (20). The precise regulatory feedback loops that detect uncontrolled proliferation and direct such cells to apoptosis are not well understood. If early cancers emerge where those mechanisms are impaired, it seemed plausible to us that assembly modulators identified through CFPSA phenotypic screens might include some chemotypes that restore the relevant feedback loops. Compounds capable of arresting proliferation, either directly or indirectly, would make potent anti-cancer agents especially if the delay in cancer progression would provide an opportunity for a patient’s innate immune system and other homeostatic feedback loops to re-establish themselves.

Abnormal signaling pathways triggered by aberrant protein-protein interactions is one way that neoplastic cells are able to achieve uncontrolled proliferation (20). In order to characterize whether assembly modulators could arrest the proliferation of neoplastic cells by redirecting key protein-protein interactions, we first sought to identify a cell line in which endogenous apoptosis was substantially lacking. While a successful anti-cancer compound would likely exhibit both anti-proliferative and cytotoxic efficacy, we wanted to conduct our screen under conditions where a readout measuring the arrest of proliferation would not be obscured by downstream activation of the normal cascade of events comprising cell death pathways.

We assessed caspase-3/7 activity in multiple tumor cell lines with an Apo-ONE assay (see **Figure 1A**). The expected correlation between endogenous triggers of apoptotic death and cancer progression was demonstrated in the LNCaP prostate cancer progression cell model, where LNCaP-C33 early (hormone sensitive) cancer cells displayed substantially more markers of apoptosis than LNCaP-C81 late (hormone resistant) cancer cells (21). In the Apo-ONE assay, CHO K1 cells show very little endogenous apoptosis (see **Figure 1A**). Hennes-20 a CHO K1 derivative into which the human APP gene has been transfected, show even less apoptosis than their parental line (see **Figure 1A**).

**Figure 1.**
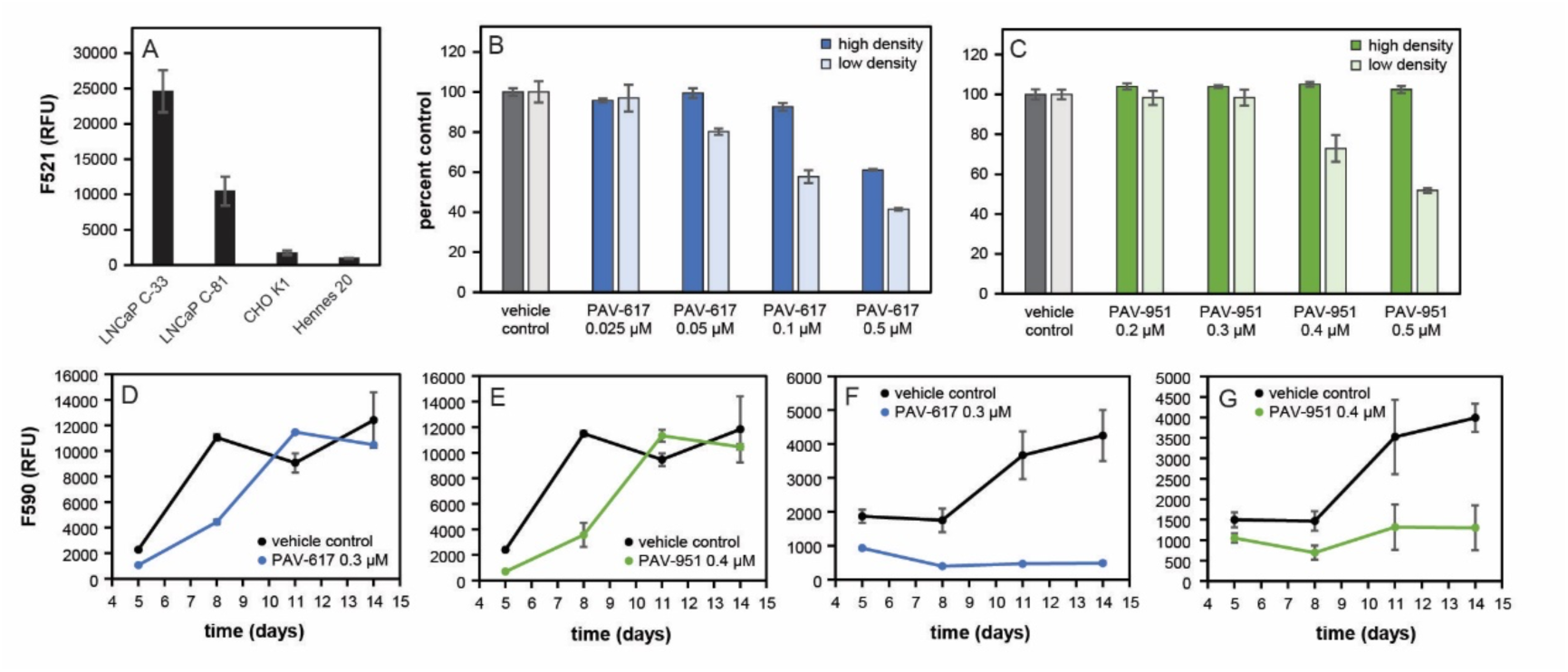
Assay development and activity of hit compounds PAV-617 and PAV-951. Figure 1A shows assessment of the endogenous apoptosis response in multiple cell lines. Plates were seeded with LNCaP C-33, LNCaP C-81, CHO K-1, and Hennes 20 cells. After three days of growth Apo-ONE reagent was added and caspase-3/7 activity was determined by fluorescent readout. Averages and standard deviation of observed activity in triplicate-repeated samples were calculated and graphed in Microsoft Excel. The Hennes 20 cell line was chosen for the counterscreen based on its low levels of caspase activity. Figures 1B and 1C show activity of PAV-617 and PAV-951 in the “arrest of proliferation” assay, where parallel plates of Hennes 20 cells were seeded at a high density of 15,000 cells per well and a low density of 500 cells per well and treated with DMSO, PAV-617, or PAV-951 in triplicate-repeated dose titrations.

Our collection of anti-viral assembly modulator compounds was then counter screened in Hennes-20 cells plated at low (500 cells/well) versus high (15,000 cells/well) densities and treated with DMSO (vehicle) or dose-titration of compounds. The rationale for this screen is than an intrinsically toxic compound should kill cells regardless of density, including in Hennes-20 cells. However, a compound that selectively triggers the arrest of proliferation will appear cytotoxic due to inhibition of cell growth when plated at a low density, but will appear non-toxic to cells plated at a high density where cells are approaching confluence and the readout detected by a cell viability assay is already close to the maximum.

Two structurally-unrelated small molecules (Tanimoto score of 41%), PAV-617 (see synthetic **Supplemental Figure 1** for chemical structure and synthetic scheme) and PAV-951 (structure not disclosed) displayed the desired phenotype (see **Figures 1B** and **1C**). The Hennes results at low versus high density comparison suggested that the effect of these compounds in cells lacking endogenous apoptosis was due to inhibition of proliferation.

To confirm that the inhibition observed in low-density Hennes 20 cells resulted from temporary arrest of proliferation and not cell death, compound was removed after a period of treatment and cell growth was measured over the subsequent two weeks for recovery potential. The cells treated with PAV-617 and PAV-951 initially showed reduced cell density relative to the control, but once compound was removed, growth was restored over time (see **Figures 1D** and **1E**). By day 11, cell density in compound-treated cells caught up to the DMSO-treated cells (see **Figures 1D** and **1E**).

Since Hennes 20 cells do not appear to have endogenous apoptosis, we sought to use a different cancer cell line to assess whether that inhibition of proliferation would be accompanied by cell death.

The recovery experiment was repeated in LNCaP C-33 cells which appear to have substantial endogenous apoptosis and treatment with PAV-617 and PAV-951 did inhibit cancer cell growth relative to the DMSO-treated control but the cells did not recover or grow significantly once compound was removed (see **Figures 1A, 1F**, and **1G**).

Fluorescent reading (RFU) corresponding to cell viability was calculated using an AlamarBlue^TM^ assay and the averages and standard deviations for triplicate-repeated samples were calculated and graphed on Microsoft Excel as a percentage of the DMSO-treated cells. PAV-617 exhibited dose-dependent inhibition of cell growth in the low density plate, indicating an EC50 between 0.1uM and 0.5uM. PAV-617 exhibited some toxicity to cells at the higher doses tested, indicating a CC50 slightly greater than 0.5uM. PAV-951 exhibited dose-dependent inhibition of cell growth in the low-density plate, indicating an EC50 around 0.5uM. PAV-951 exhibited no significant inhibition of cell growth at the tested doses in the high-density plate, indicating a CC50 greater than 0.5uM. **Figures 1D**-**1G** show recovery of cancer cell growth following removal of compound. Hennes 20 or LNCaP C-33 cells were seeded at a low density then incubated with DMSO, PAV-617, or PAV-951. After a period of treatment, the medium containing compound was removed and replaced with fresh media. Plates were assessed for cell viability by AlamarBlue^TM^ on days 5, 8, 11, and 14 and the averages and standard deviations of triplicate-repeat samples were calculated and graphed over time on Microsoft Excel. PAV-617 and PAV-951 treated Hennes 20 and LNCaP C-33 cells all showed reduced viability compared to matched DMSO-treated cells on day 5. However, cell growth in the Hennes 20 cells which had been treated with compound recovered with time (**Figure 1D, E**), while the LNCaP C-33 cells which had been treated with compound did not recover (**Figure 1F, G**).

### Investigating the Activities of PAV-617 and PAV-951: from modulators of viral capsid assembly to pan-cancer therapeutics

The anti-proliferative compounds PAV-617 and PAV-951 had originally emerged from our CFPSA screen as inhibitors of viral capsid formation. The CFPSA model had been validated by demonstrating that antiviral assembly modulator hits display activity against infectious viruses in cell culture (9,11–13). PAV-617 is active against pox viruses in cell culture (13). The effective concentration for half maximal activity (EC50) of PAV-617 against monkeypox (MPXV) is approximately 300 nM (see **Figure 2A**). PAV-951 is active against HIV in cell culture with an EC50 between approximately 300 nM and 1uM (see **Figure 2B**).

**Figure 2.**
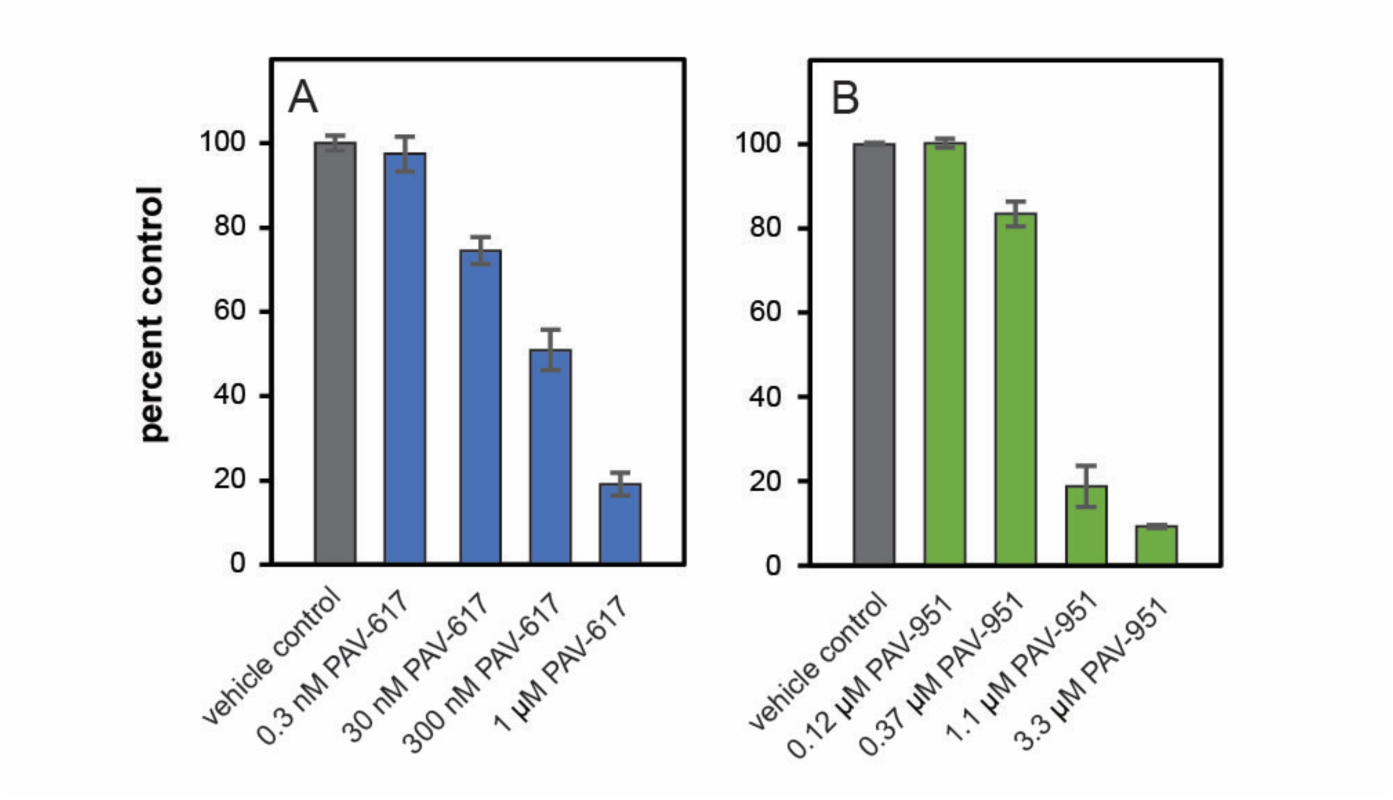
Anti-viral properties of PAV-617 and PAV-951. Figure 2A shows activity of PAV-617 against MPXV. BSC-40 cells were infected with 100 plaque forming units of MPXV Zaire 79 and treated with PAV-617 for three days. Averages and standard deviation for plaques observed with triplicate-repeated dose-titration of PAV-617 are shown as a percentage of the plaques observed in untreated cells with an EC50 of approximately 300 nM. Figure 2B shows activity of PAV-951 against HIV. MT-2 cells were infected with NL4-3 Rluc HIV and treated with PAV-951 for four days. Averages and standard deviation of viral titer observed with triplicate-repeated dose-titration of PAV-951 are shown as a percentage of the titer observed in DMSO-treated cells, with an EC50 between 0.37 uM and 1.1 uM.

When PAV-617 and PAV-951 were identified as having anti-cancer activity in addition to anti-viral properties, we suspected that the compounds might correct cancer-induced defects in protein assembly that were related to the aberrant assembly induced by viruses to promote capsid assembly. To get a better understanding of how the defects present themselves across diverse cancers, we assessed the anti-cancer activity of these two compound chemotypes on a panel of 15 cancer cell lines from the Eurofins OncoPanel^TM^. The cell lines were derived from a variety of tissues and were representative of male and female patients of different ages from pediatric to senior. Both PAV-617 and PAV-951 showed pan-cancer activity with dose-dependent tumor growth inhibition in all 15 cell lines (see **Figures 3A** and **3B**).

**Figure 3.**
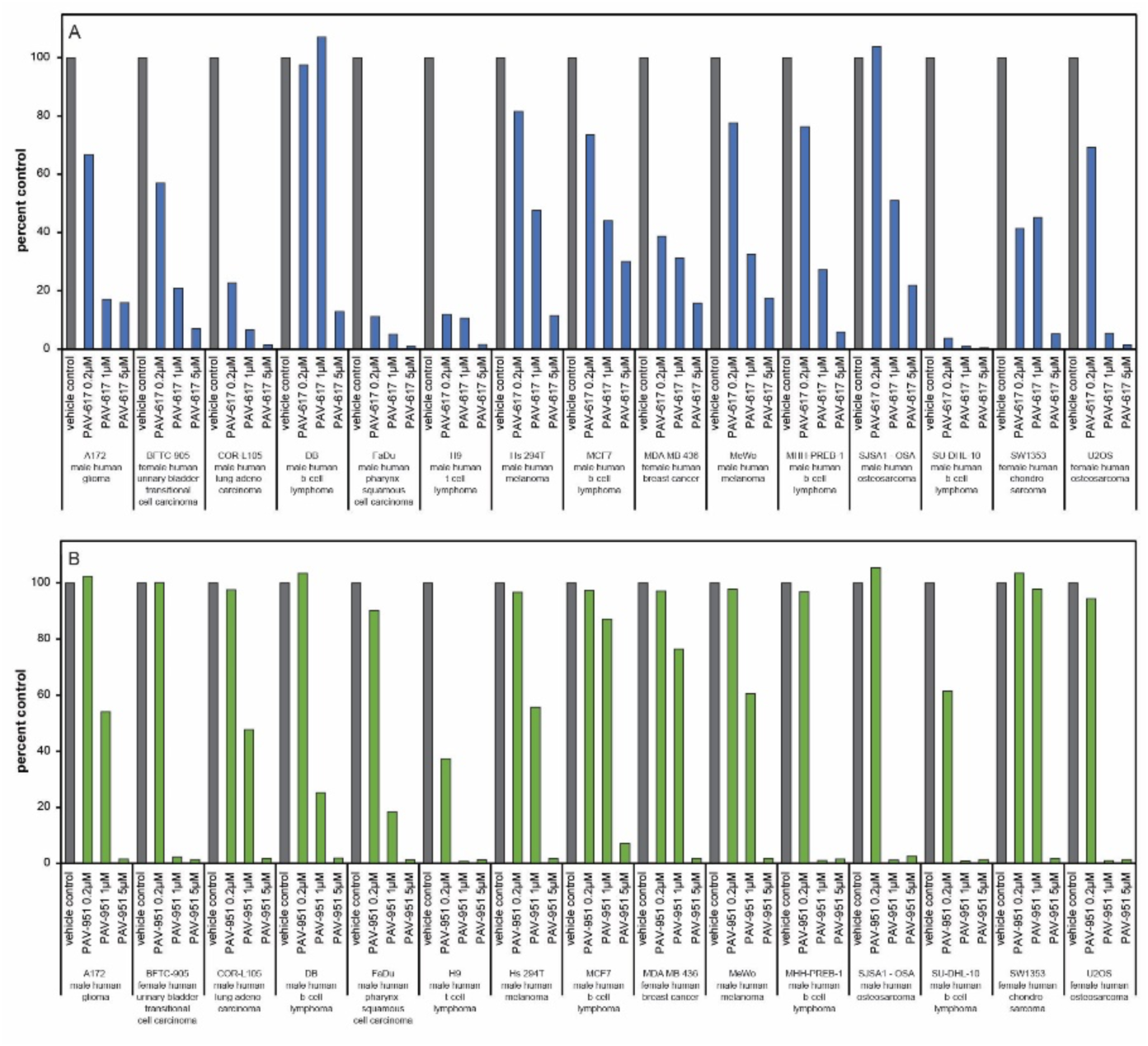
Pan-cancer activity of PAV-617 and PAV-951. **Figures. 3A** and **3B** show results from a panel of human tumor cell lines— A172 (male human glioma), BFTC-905 (female human urinary bladder transitional cell carcinoma), COR-L105 (male human lung adenocarcinoma), DB (male human b-cell lymphoma), FaDu (male human pharynx squamous cell carcinoma), H9 (male human t-cell lymphoma), Hs 294T (male human melanoma), MCF7 (female human breast cancer), MDA MB 436 (female human breast cancer), MeWo (male human melanoma), MHH-PREB-1 (male human b-cell lymphoma), SJSA1-OSA (male human osteosarcoma), SU-DHL-10 (male human b-cell lymphoma), SW1353 (female human chondrosarcoma), and U-2 OS (female human osteosarcoma). Cells were grown for 24 hours then treated with either vehicle, PAV-617, or PAV-951. Cell viability after 3 days of treatment was measured as bioluminescence intensity and averages of triplicate-repeat dose-titrations with PAV-617 and PAV-951 were graphed on Microsoft Excel as percent of bioluminescence observed in DMSO-treated cells. Both compounds showed inhibitory effects in all 15 cell lines.

Confirmation of the pan-cancer activity of these compounds was received through a sixty cancer line screen carried out by the National Cancer Institute (see **Supplemental Figure 2**)

### Animal validation of PAV-617 and PAV-951 anti-cancer efficacy

With demonstrated anti-viral activity, demonstrated anti-cancer activity, and data supporting a proliferation-based mechanism of action, we assessed mouse toxicology and pharmacokinetic (PK) properties in order to determine suitability for efficacy studies in an animal model.

The maximum tolerated dose (MTD) estimates how much compound can be administered to an animal without adverse effects (22). When administered orally to mice, both PAV-617 and PAV-951 displayed MTDs greater than 20 mg/kg, which was the highest dose tested, as no clinical symptoms or significant differences were observed between vehicle and treatment groups. When administered by intraperitoneal (IP) injection, the MTD of PAV-617 was safe at 10 mg/kg. When administered by IP injection, the MTD of PAV-951 was determined to be 2.5 mg/kg. Subsequently, chemical analogs of PAV-951 have been identified with MTDs of 15 mg/kg when dosed IP. Some of those analogs, including compound PAV-442, displayed comparable activity as PAV-951 against A549 lung cancer and PANC-1 pancreatic cancer tumor lines in cell culture (see **Supplemental Figures 3A** and **B** for activity of PAV-442). PAV-181, a chemical analog of PAV-617, was found to be safe in mice at 20 mg/kg (higher doses were not tested). PAV-181 retained the pan-cancer activity of PAV-617 in the Eurofins OncoPanel^TM^ (see **Supplemental Figure 3C** for the disclosed chemical structure of PAV-181 and **3D** for anti-cancer activity).

The PK properties are based on the absorption, distribution, metabolism, and excretion of a compound in a living organism and are necessary to determine dosing parameters because any compound designed for clinical use needs to achieve an efficacious concentration in a target organ (23,24). As an early measure of PK, we determined the concentration of compound in the plasma of rats or mice over time following one intravenous (IV), one IP, or one oral dose. Both compounds were detectable in the animals through all administration routes, though the maximum concentration achieved (Cmax) and rate of elimination were variable across conditions (See **Supplemental Figure 3**).

We determined that, while both chemical series would need optimization on the PK and toxicological properties before being named as clinical drug-candidates, PAV-617 and PAV-951 would be adequate for a preliminary animal efficacy study in order to validate whether or not the anti-proliferative properties of the compounds observed in cell culture translates to anti-proliferative properties in animals.

In the first set of animal efficacy studies, human A549 non-small cell lung cancer cells were grafted subcutaneously onto mice. After 30 days of tumor establishment, the animals received daily treatment with PAV-617 or PAV-951 and tumor volume was measured over time. The doses and routes of administration for PAV-617 (10 mg/kg IP injection) and PAV-951 (1.5 mg/kg IV injection) were determined based on their MTD and Pk properties. The PAV-617 study was conducted for 28 days and the PAV-951 study was conducted for 14 days. As negative and positive controls, both studies included a group treated with vehicle only and a group treated with Gemcitabine hydrochloride, an FDA approved drug for non-small cell lung cancer, administered to animals in the same way as the test compound. The Gemcitabine was administered at the standard dose of 100 mg/kg. Both PAV-617 and PAV-951 reduced tumor growth significantly compared to the vehicle-only groups and performed comparably to Gemcitabine despite being administered at substantially lower doses. PAV-617 displayed a tumor growth inhibition (TGI) of 63% compared to Gemcitabine’s TGI of 64% while PAV-951 had a TGI of 72% compared to Gemcitabine’s TGI of 84% (see **Figures 4A** and **4B**).

**Figure 4.**
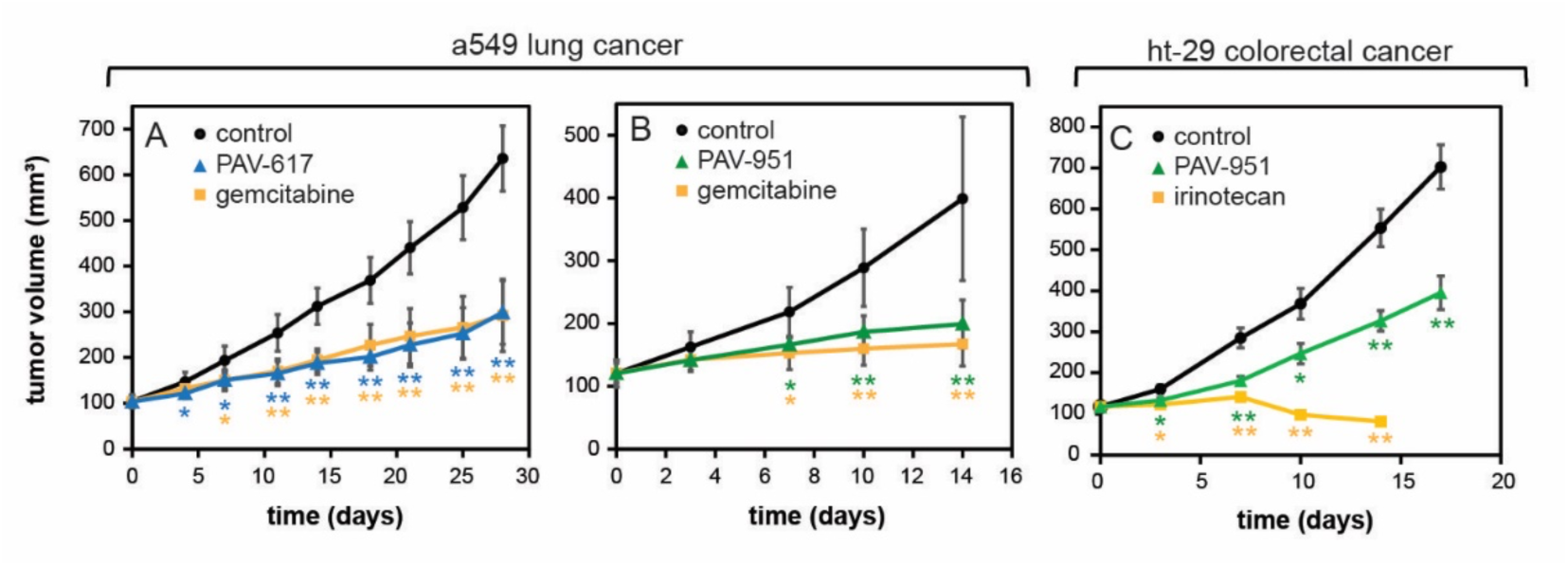
Antiproliferative activities of PAV-617 and PAV-951 in animal cancer xenografts. Tumors were grafted onto mice via a subcutaneous injection and grown until they reached a volume of 100mm^3^. Animals were divided into randomized groups and treated with vehicle, PAV-617, PAV-951, or an FDA approved cancer drug as a positive control. Figure 4A shows tumor volume across time for an A549 lung cancer treated with vehicle, PAV-617, or Gemcitabine hydrocholoride. Figure 4B shows tumor volume across time for a A549 lung cancer treated with vehicle, PAV-951, or Gemcitabine hydrochloride. Figure 4C shows tumor volume across time for a HT-29 colorectal cancer treated with vehicle, PAV-951, or Irinotecan.

In the second animal efficacy study, human HT-29 colorectal adenocarcinoma cells were grafted subcutaneously onto mice. After tumor establishment, the animals received treatment with vehicle, 3 mg/kg PAV-951, or 60 mg/kg of a positive control drug Irinotecan. After 17 days of treatment, PAV-951 had significantly reduced tumor growth relative to the vehicle-only group (TGI of 52%), though the impact was lower than Irinotecan (TGI of 108%) (See **Figure 4C**).

### Characterizing the targets of PAV-617 and PAV-951

As PAV-617 and PAV-951 were identified by a phenotypic screen, their actual targets were unknown during the early stages of compound advancement. To identify their targets, each molecule was coupled to Affi-gel resins from a position on the molecule unrelated to proliferation arrest activity based on structure-activity-relationship (SAR) exploration (see **Supplemental Figure 1B** for the synthetic scheme of the PAV-617 resin used). In that way, they could serve as target-binding ligands for drug resin affinity chromatography (DRAC) (25). LNCaP C-33 cells were chosen for DRAC starting material because we knew that PAV-617 and PAV-951 displayed efficacy against them and we wanted a model for our target engagement studies that would account for mechanisms of endogenous apoptosis responses. Extracts were prepared from LNCaP C-33 cells that were treated for 22 hours with either DMSO, PAV-617, or PAV-951. The extracts were applied to the PAV-617, PAV-951, or control drug resins, washed with 100 bed volumes of buffer, and eluted with either 100 uM PAV-617 or 100 uM PAV-951.

Samples of the DRAC eluate were sent for analysis by tandem mass spectrometry (MS-MS). The DRAC eluate from the PAV-617 and PAV-951 resins contained large sets of proteins missing from the control resin eluate. This included cancer-implicated proteins from the literature. 92 proteins from the DMSO-treated cell extracts were identified in the PAV-617 resin eluate that were not present in the control resin eluate. 116 proteins from the DMSO-treated cell extracts were identified in the PAV-951 resin eluate which were not present in the control resin eluate. Of the proteins identified by MS-MS as unique or greatly enriched in the drug resin eluates, 38 proteins in the DMSO-treated cell extracts were unique to PAV-617, 62 proteins in the DMSO-treated cell extracts were unique to PAV-951, and 54 proteins from the DMSO-treated cell extracts were found in both the PAV-617 and the PAV-951 resin eluates (see **Figures 5A-C**). When the proteins detected in the eluates were searched in a database for cancer-implicated proteins, 23 of the proteins from the PAV-617 resin eluate and 29 proteins from the PAV-951 resin eluate were known to be part of cancer-relevant interactomes (See **Figures 5A-C**).

**Figure 5.**
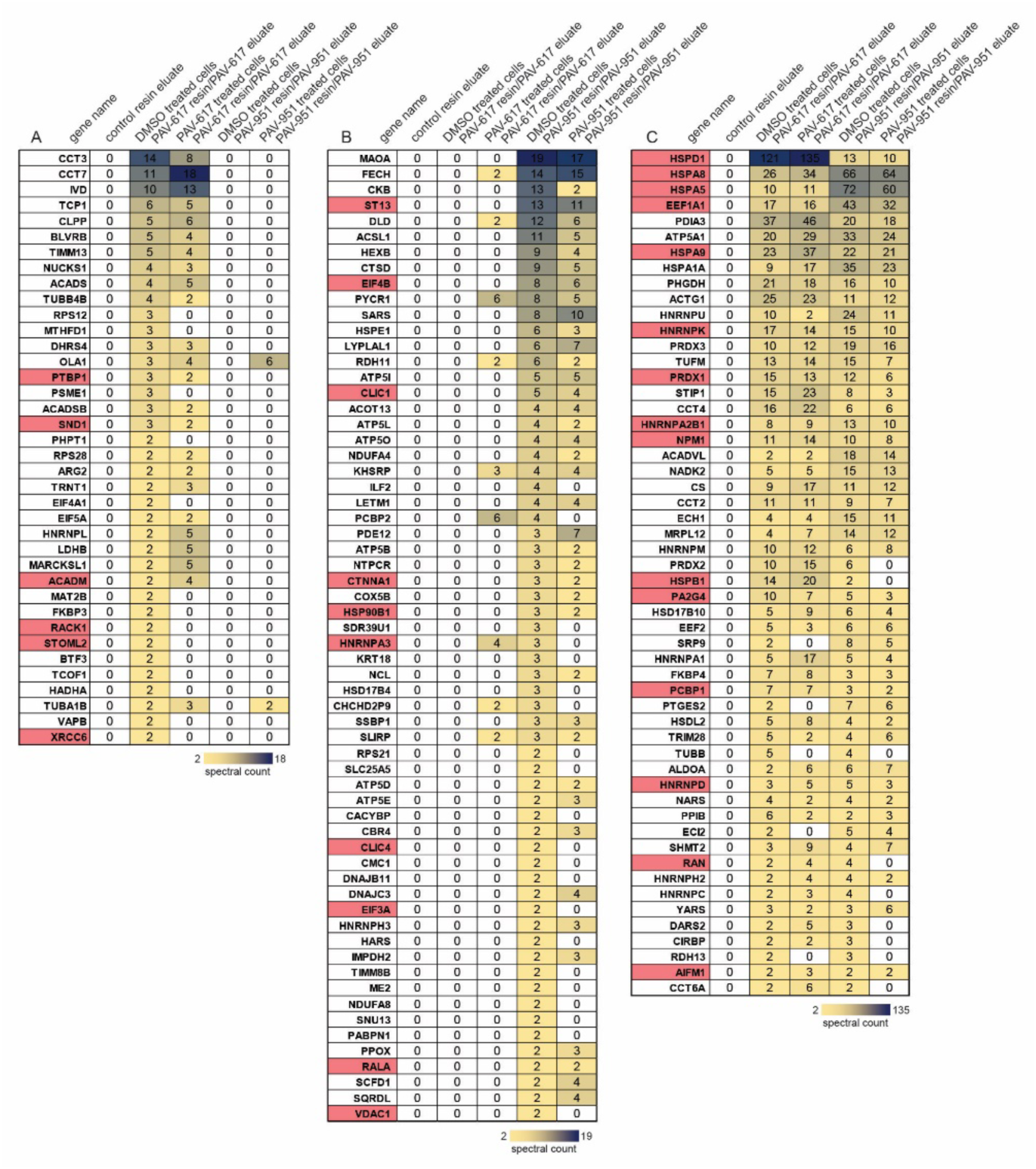
Identification of proteins in PAV-617 and PAV-951 resin eluates by MS-MS. DRAC experiments were performed where 30 ul of DMSO or compound-treated LNCaP cell extract, adjusted to a protein concentration of approximately 10 mg/ml in column buffer, was incubated on a column containing 30ul of affigel resin coupled to either PAV-617, PAV-951, or a 4% agarose matrix (control) for one hour at 4 degrees Celsius. The input material flow-through was collected and the resin was washed with 3 mL column buffer then eluted overnight at 4 degrees Celsius in 100 ul of either 100uM PAV-617 or 100uM PAV-951 in column buffer. Figures 5A**-5C** show spectral counts of proteins detected by MSMS in single-point DRAC eluates. Figure 5A shows the set of proteins only detected in PAV-617 resin eluate. Figure 5B shows the set of proteins only detected in the PAV-951 resin eluate. Figure 5C shows the set of proteins detected in both the PAV-617 and the PAV-951 resin eluates. Conditional formatting has been applied where relative abundance (by spectral count) of a given protein in a sample is visualized on a yellow-to-black scale. Proteins implicated with cancer in the Bushman labs oncogene database (http://www.bushmanlab.org/links/genelists) have been indicated in red.

The MS-MS indicated that when LNCaP C-33 cells were treated with compound, composition of the eluate was subsequently affected. For both the PAV-617 and the PAV-951 resin eluates, the spectral counts of some particular proteins detected by MS-MS increased or decreased in treatment conditions (See **Figures 5A-C).** However, for other proteins, the number of spectral counts detected in the eluates remained unchanged upon treatment. When we entered the proteins identified from the DRAC eluate into a database of protein-protein interactions, we discovered that many of the proteins eluted by either or both PAV-617 or PAV-951 were known to be involved in networks that involved frequent interactions and associations with each other (see **Supplemental Figure 5**) and relevance for cancer (see **Supplemental Figure 6**).

DRAC experiments were conducted side-by-side in triplicate with and without addition of metabolic energy by running the experiment at 4°C versus 22°C and supplementing with an “energy cocktail” of ribonucleotide triphosphates (1mM rATP, 1mM rGTP, 1mM rCTP, 1mM UTP) and 5ug/mL creatine kinase. Changes in the amounts of proteins that bound and eluted with PAV-617 or PAV-951 were observed by western blot in the presence of these metabolic energy substrates (see **Figures 6A** and **6B**). For the PAV-617 resin eluate, B-tubulin (TUBB), KRAB-associated protein 1 (KAP1 or TRIM28), Methylenetetrahydrofolate dehydrogenase 1 (MTFGD1), and protein disulfide isomerase (PDI)—proteins which had previously been identified in the eluate by MS-MS—all showed up by western blot in larger amounts under energy-supplemented conditions (See **Figure 6A**). For the PAV-951 resin eluate, MTHFD1, PDI and heterogenous ribonucleoprotein K (hnRNPK) were observed to be enhanced under energy-supplemented conditions (see **Figure 6B**). By contrast, the amount of KAP1 identified in the PAV-951 resin eluate decreased in the presence of metabolic energy substrates (see **Figure 6B**).

**Figure 6.**
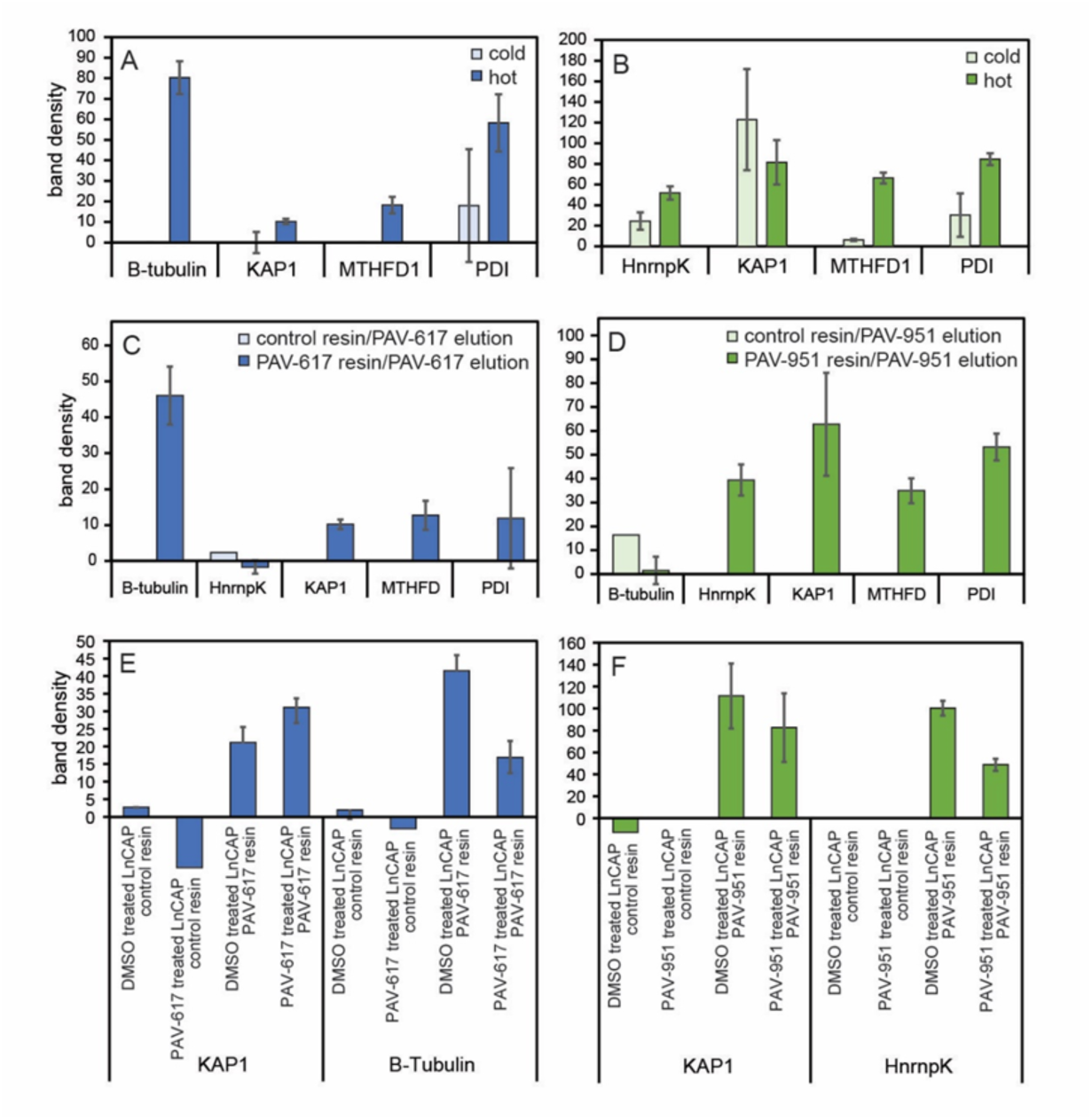
Exploration of changes to PAV-617 and PAV-951 resin eluates under different conditions. DRAC experiments using LnCAP C-33 starting extract as described in **Fig. 5** were conducted under conditions which included supplementing the starting material and eluate with an energy cocktail to a final concentration of 1mM rATP, 1mM rGTP, 1mM rCTP, 1mM UTP, and 0.05 ug/mL creatine kinase and running the experiment at room temperature versus cold conditions at 4 degrees Celsius and without the energy cocktail. **Figures 6A** and **6B** show quantitation of average integrated density of protein band detected by western blot in triplicate-repeat samples eluted with PAV-617 from the PAV-617 or eluted with PAV-951 from the PAV-951 resin when DRAC was conducted side-by-side under energy-supplemented warm and non-energy-supplemented cold conditions. **Figures 6C** and **6D** show quantitation of average integrated density of protein band detected by western blot for TUBB, hnRNPK, KAP1, MTHFD1, and PDI in triplicate-repeat eluates under energy-supplemented conditions from the PAV-617 resin or PAV-951 resin and a single point elution with each compound from the control resin. Resins were eluted with either compound or 1% DMSO and the amount of protein detected in the 1% DMSO elution is subtracted from the compound elution in the figure. **Figure 6E** shows quantitation of average integrated density of protein band detected by western blot for KAP1 and B-tubulin in DRAC eluates generated under energy-supplemented warm conditions of DMSO versus PAV-617 treated cells eluted with PAV-617 in triplicate from the PAV-617 resin and in single-point from the control resin. **Figure 6F** shows quantitation of average integrated density of protein band detected by western blot for KAP1 and hnRNPK in DRAC eluates generated under energy-supplemented warm conditions for DMSO versus PAV-951 treated cells eluted with PAV-951 in triplicate from the PAV-951 resin or in single-point from the control resin.

To confirm that the enrichment required the combination of metabolic energy substrates and compound rather than the presence of energy substrates alone, resins were eluted side-by-side in triplicate with either PAV-617 or PAV-951, or 1% DMSO all containing the energy cocktail. By western blot, eluates from both PAV-617 and PAV-951 resins were found to contain significant amounts of TRIM28/KAP1, MTHFD1, and PDI relative to the control resin or DMSO elution. The eluate of the PAV-617 resin contained TUBB which the PAV-951 resin eluate did not have in any greater amount than the control. The eluate of the PAV-951 resin contained hnRNPK, not present in the PAV-617 resin eluate in greater amount than the control.

DRAC experiments were also carried out under conditions where the starting extract was prepared from LNCaP C-33 cells that had been treated with DMSO, 100 uM PAV-617, or 100 uM PAV-951 for 24 hours, in order to see if the changes to the eluate observed by MS-MS would repeat with energy supplementation. Western blots of triplicate-repeated samples showed the PAV-617 resin eluate contained increased amounts of KAP1 and decreased amounts of TUBB when cells were treated with PAV-617, meanwhile PAV-951 resin eluate showed decreased amounts of both KAP1 and hnRNPK when cells were treated with PAV-951 (See **Figures 6E-F**).

One explanation for why the DRAC eluates contain large numbers of proteins is that the targets for PAV-617 and PAV-951 are themselves multi-protein complexes, as had been observed for analogous studies on structurally unrelated protein assembly modulator chemical series effective in other therapeutic areas. To test this hypothesis and determine which proteins directly bind the compounds and which are indirectly associated with the compounds via protein-protein interactions involving the direct drug-binding protein(s), we modified the compounds into photocrosslinker analogs by attachment of diazirine and biotin functional groups at the same position to where the resin had previously been attached. The photocrosslinker analogs were designed so that after an incubation with cell extract that would allow the compound to bind its target, with subsequent exposure to ultraviolet light forming a covalent bond between the diazirine moiety of the compound and the nearest protein (26). The sample could then be solubilized and precipitated with streptavidin beads (which bind biotin) to identify the drug-binding protein(s). The streptavidin precipitation (SAP) could be done using a native sample, which would pick up the direct drug-binding protein(s) and with it any co-associated proteins. Or the SAP could be done using a denatured sample which would, by virtue of the covalent bond to biotin, identify only the direct drug binding protein(s), with all other associated proteins of the target multi-protein complex lost upon denaturation.

A549 cell extract was incubated with either 1% DMSO or the photocrosslinker analogs of PAV-617 or PAV-951, then exposed to ultraviolet light. The samples were then divided into two equal parts, where one part was left native and the other denatured, then both were adjusted to non-denaturing conditions and incubated with streptavidin beads. Blots of the SAP samples for KAP1 indicate KAP1 is only a component of the PAV-617 target under native conditions and is completely lost upon denaturation (See **Figures 7A-B**). However, KAP1 is present to a significant extent in both native and denatured conditions for the SAP with the PAV-951 crosslinker (see **Figures 7A** and **7C**). The SAP samples were sent for MS-MS analysis, however high background in the samples with no crosslinker added rendered the data uninformative for definitive binding partner identification (data not shown).

**Figure 7.**
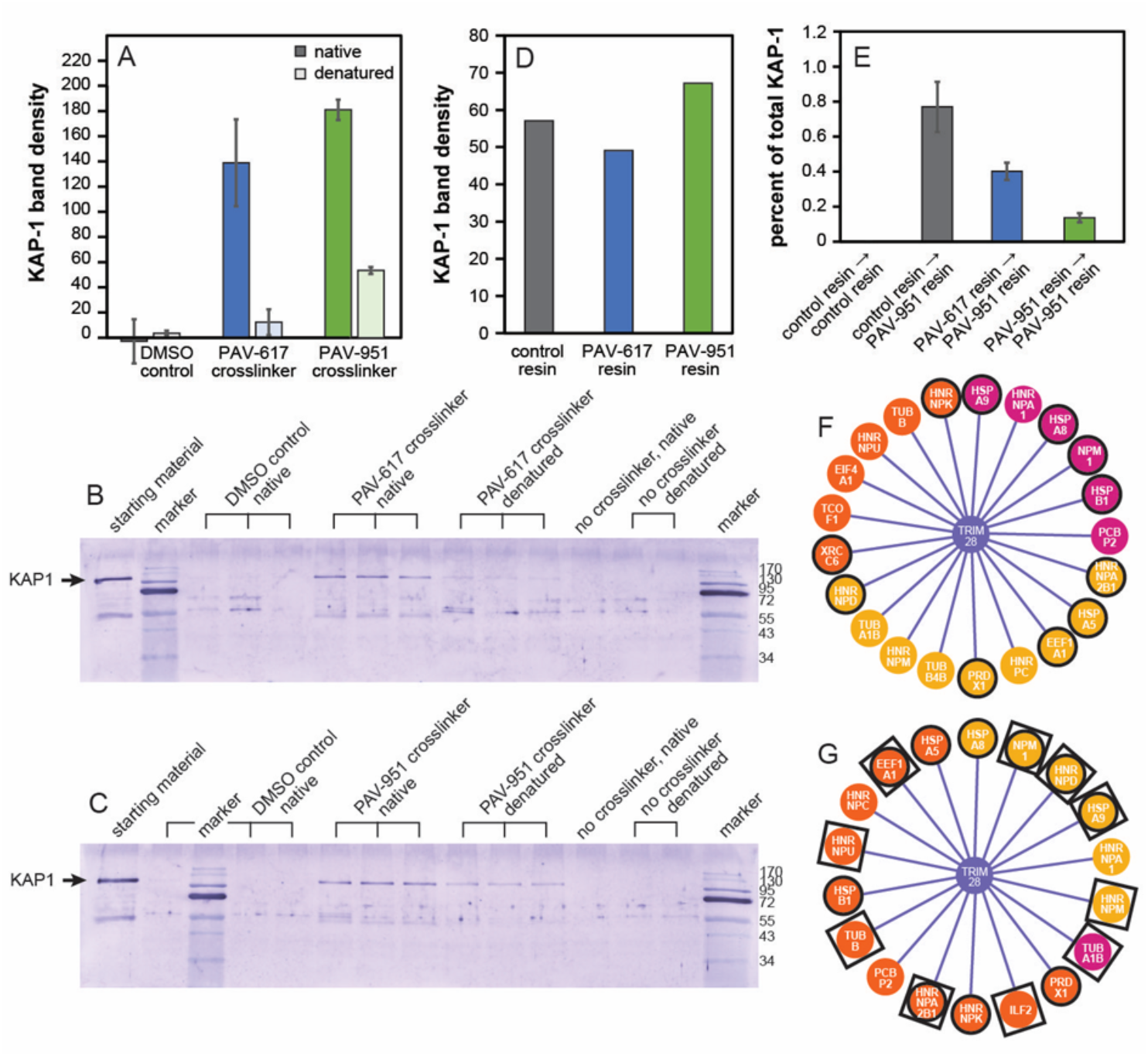
A subfraction of KAP-1/TRIM28 is a component of the PAV-617 and PAV-951 target complexes. Figure 7A shows quantitation of average integrated density of KAP1 protein band detected by western blot while Figures 7B and 7C show the western blots themselves, in triplicate-run native and denatured streptavidin precipitations of photo crosslinked samples. Crosslinking experiments were performed where 65 uL of A549 cell extract was adjusted to a protein concentration of approximately 1 mg/ml in column buffer and supplemented with the energy cocktail, with either 1% DMSO or 1uM modified photo crosslinker analogs of PAV-617 and PAV-951 for one hour at room temperature then 20 minutes on ice. The extracts were exposed to UV light then divided into two aliquots. One aliquot was left native and one was denatured by adding DTT, SDS, and boiling. 800 uL of column buffer with 0.1% triton was added to both aliquots and then they were incubated with 2.5ul magnetic streptavidin beads for one hour at room temperature before being denatured in loading buffer containing SDS and heated to 100°C for 3 minutes. Figures 7D and 7E show the show quantitation of integrated density of KAP1 protein band detected by western blot in LNCaP extract depleted on the PAV-617, PAV-951, and control resins. 230 uL of LNCaP extract was incubated with 230uL of PAV-617, PAV-951, or control resins in single point for one hour in energy-supplemented conditions. Depleted flow-throughs were divided and put onto a subsequent PAV-951 column in triplicate or onto a subsequent control column in single-point under energy-supplemented conditions. Columns were eluted three times-a first overnight elution with PAV-951, a second overnight elution with PAV-951, and a third elution with 1% SDS. Eluates were diluted 3:1 in loading buffer and analyzed by western blot. Every western blot for the eluate included a sample of the original, un-depleted starting LNCaP extract diluted 1:100 in loading buffer. Figure 7D shows quantitation from when the flow-throughs from each columns were blotted for KAP1 to determine how much KAP1 had been depleted from each resin. Figure 7E shows the amount of protein detected in the eluate normalized as percent of their corresponding total sample, where the amount detected by eluate samples were divided by the amount detected in the starting material sample, then multiplied by 0.013 to match the concentrations. The percent detected by western blot from the two overnight elutions and the SDS elutions were added together to determine the total percentage of cellular KAP1 was binding to and eluting from the resins). Figures 7F and 7G show diagrams of proteins identified by MS-MS in Figs. 5A-5C as comprising the PAV-617 resin/eluate and PAV-951 resin/eluate found in the NURSA database of protein-protein interactions as interacting with KAP1 (https://dknet.org/about/NURSA_Archive)(27). Orange indicates KAP1 implicated proteins detected in the eluate which decreased with treatment. Magenta indicates KAP1 implicated proteins detected in the eluate which increased with treatment. Yellow shows KAP1 implicated proteins detected in the eluate which were unchanged with drug treatment. Circles indicate proteins from the PAV-617 and PAV-951 resin eluates implicated in cancer from the Bushman lab oncogene database (http://www.bushmanlab.org/links/genelists). Squares indicate proteins from the PAV-951 resin eluates implicated in HIV from the virus mentha database (https://virusmentha.uniroma2.it/) (28).

Conventional methods of drug discovery typically involve the use of recombinant proteins to measure affinity between a drug and its target (23). However, we have previously observed that protein assembly modulating compounds are specific to their target proteins only when that protein is found in particular cellular contexts/co-associations and we were concerned that if the protein-protein interactions between the direct drug-binding protein and other proteins comprises an important dimension of PAV-617 and PAV-951’s targets, isolated recombinant proteins would not be an appropriate surrogate for the protein-protein interactions occurring *in vivo*. To measure target engagement for PAV-617 and PAV-951, we returned to DRAC and determined whether passing cell extract over the PAV-617 or PAV-951 resins would deplete the extracts of bindable target.

We applied the flow-through of PAV-617, PAV-951, and control resins to a second copy of the drug resin for these reasons and demonstrated that the drug resins deplete the extract of essentially all KAP1 capable of binding to the resins (see **Figures 7D-E**). Western blot of the resin flow-through showed that a comparable amount of KAP1 was flowing through the PAV-617, PAV-951, and control resins without binding. However, when the flow-through that had been depleted on the PAV-951 resin was applied on to a new PAV-951 resin, very little additional KAP1 bound to the second resin (see **Figure 7E**). By contrast, application of the control resin flow-through (which was not specifically depleted of anything) to a second PAV-951 resin, showed significant KAP1 binding to the second resin. Application of the PAV-617 resin flow-through to the PAV-951 resin showed significantly more KAP1 binding to the second resin than from the PAV-951 flow through, but significantly less KAP1 binding than from the control flow through (see **Figure 7E**).

One notable observation was that, even though the five-fold depletion by the PAV-951 resin of its target compared to the control resin was statistically significant and reproducible, it accounted for a tiny amount of the total KAP1 which had been detected by western blot in the starting extract. Only 0.7% of the total amount of KAP1 detected in the starting material bound to the PAV-951 resin after passing over a control column (see **Figure 7E**). As a reference point to account for non-specific loss of material of the course of the experiment (due to denaturation over time or nonspecific sticking removed during the washing phase)—of the original PAV-951 resin with which the extract was depleted, serial elution with PAV-951 and 1% SDS showed approximately 4% of the total KAP1 detected in the original extract had bound to the resin (data not shown). We conclude that the compounds selectively target this small subfraction of KAP1 (less than 5% of the total KAP1 in the extract) because once the subfraction of KAP1 capable of binding to the resins is removed, the remaining extract will not bind anymore, even though it still contains ample KAP1.

Several of the proteins found by MS-MS in the PAV-617 and PAV-951 resin eluates are known to interact directly with KAP1 (see **Figures 7F** and **7G**). These KAP1 implicated proteins include some whose relative amount in the eluate increased and/or decreased with drug treatment and are part of cancer associated interactomes identified from the literature (see **Figures 7F** and **7G**). The KAP1 implicated proteins from the PAV-951 resin/eluate also contained several proteins that are associated with HIV in the literature (see **Figure 7G**). These associations may shed light on the cellular role played by the subfractions of KAP1 that are targeted by PAV-617 and PAV-951.

## Discussion

Our data indicate that PAV-617 and PAV-951, two protein assembly modulator anti-viral compounds selected for their ability to arrest proliferation in a distinctive way by a novel screen, are cytotoxic to a wide range of neoplastic cell lines representing both rare and common cancers. These compounds, while early in their drug optimization, performed comparably to the commercial anti-cancer drug Gemcitabine, at 10 and 60 fold lower doses, to inhibit the growth of an A549 tumor in immunodeficient mice. PAV-951 was also able to reduce growth of a HT-29 colorectal cancer in a mouse xenograft study, though to a lesser degree than the positive control Irinotecan. These data suggest that these two compounds are directed to two different targets common to a wide range of cancers, and may provide a starting point for the development of novel cancer therapeutics.

DRAC and photocrosslinking experiments indicate that PAV-617 and PAV-951 interact with proteins that are part of a multi-protein complex. These complexes are dynamic, as demonstrated by changes in the eluate when DRAC is carried out in the presence versus absence of metabolic energy substrates or from untreated versus compound-treated cell starting extract. Together, these findings suggest a new model for disease pathogenesis in which previously unappreciated transient multi-protein complexes plays an important role in the dynamics linking, in this case, cellular proliferation to apoptosis. We hypothesize that cancer progression is facilitated by aberrant versions of these multiprotein complexes in which the linkage of inappropriate proliferation to apoptosis is attenuated or slowed. The effect of protein assembly modulating drugs is to restore the original version of the multi-protein complex, possibly through an allosteric mechanism-of-action (29).

KAP1/TRIM28, an identified protein component of the PAV-617 and PAV-951target complexes, is worthy of specific mention. KAP1 was identified by both MS-MS and western blot as being part of the PAV-617 and PAV-951 targets (see **Figures 5-7**). Crosslinking experiments showed that for PAV-617 KAP1 is present in a complex targeted by the compound under native conditions, but is lost upon denaturation, indicating it is not the direct drug-binding protein, but rather more likely a distal component of the target multi-protein complex (see **Figure 7**). For PAV-951, KAP1 is also part of the complex under native conditions. However the data indicates that a portion of KAP1 is a direct drug-binding protein as well. This suggests more than one copy of KAP1 per multi-protein complex (see **Figure 7**), and is consistent with the hypothesis that KAP1 has multiple functions.

KAP1 stood out as being of particular interest because it is a known allosteric modulator, implicated in both infectious and noninfectious disease (30). KAP1 is involved in a variety of protein-protein interactions and an array of functions including transcriptional activation of HIV, T-cell development, DNA damage repair, as a transcriptional co-repressor for many genes, and as a ligase for post-translational modifications such as ubiquitination and SUMOlyation (30,31). Studies have shown increased levels of KAP1 in many types of cancer and high levels of KAP-1 correlate with aggressive clinical phenotype and progression to metastasis (32–34). However, other described functions of KAP1 are tumor suppressive and promote autophagy (30,35,36). KAP1 directly binds with other cancer-implicated proteins including MDM2, TRIM14, Fructose-1,6-biphosphatase (FBP1), MAGE-A3, MAGE-C2, Heat shock protein 70, and TWIST1 (36–40). The diversity of functions that KAP1 displays appears to be, at least in part, through its assembly into different multi-protein complexes that carry out different objectives.

The literature describes that KAP1 is utilized by HIV to the host’s detriment but also functions as a key component of the host’s immune response in repressing HIV (30,41). KAP1 is implicated as a key part of a pathway hijacked by the poxvirus p28 virulence factor (42). We show that PAV-951 is active against HIV and PAV-617 is active against MPXV in addition to their anti-cancer phenotypes (see **Figure 2**). We hypothesize that there are shared alterations in protein assembly in neoplastic cells and virus-infected cells. Advancing the SAR of PAV-617 and PAV-951 could be utilized to determine whether the antiviral and anti-cancer targets are identical or merely similar. In the latter case, the activities would separate with further SAR. Regardless, these findings suggest that the initial CFPSA capsid assembly screen was successful in identifying novel targets relevant to both viruses and cancer.

We suggest that PAV-617 and PAV-951 redirect protein-protein interactions, whereby some specific proteins are recruited to, and other specific proteins are expelled from, distinct multi-protein complexes in the presence of compound (see **Supplemental Figure 7** for a diagram). KAP1 is known to both promote and suppress tumorigenesis (30). Therefore, a compound which selectively targets some forms of KAP1 and not others would be important regardless of the nature of the interaction with KAP1. In some cases, as observed for PAV-951, a portion of KAP1 is a direct drug-binding protein, although other copies of KAP1 appear not to be. In the case of PAV-617 resin, KAP1 is associated with the drug only indirectly by virtue of being a protein present in the target multi-protein complex, but at a distance from the drug-binding site, thus present by SAP under native but not denatured conditions.

We interpret the DRAC flow-through data to mean that, by virtue of conformation or other differences among co-associated proteins in a transient multi-protein complex, subfractions of cellular KAP1 are selectively targeted by PAV-617 and PAV-951. Results from the cross-depletion where the flow-through of the PAV-617 resin was applied to a new PAV-951 resin further indicate that the PAV-617-binding subfraction of KAP1 must be distinguishable from the PAV-951 binding subfraction of KAP1 because depletion of one does not fully deplete the other (see **Figure 7E**).

These findings about PAV-617 and PAV-951 mirror those made for PAV-431, a structurally-unrelated protein assembly modulator with pan-respiratory antiviral activity (9). It appears as though protein assembly modulating compounds, despite structural diversity, share characteristics including the targeting of multi-protein complexes, selectivity for a small fraction of the total amount of a given protein found in a cell, and allosteric mechanisms of action (9,11,29). Furthermore, they appear to have remarkable activity across broader categories of pathogens (e.g. pan-viral family and pan-cancer) than is generally observed for existing drugs. The antiviral assembly modulators appear to have a barrier to the development of resistance (9,12). Further studies are needed to determine whether this property holds true for the anti-cancer subset of protein assembly modulators.

Effective cancer drugs have been developed based on a number of mechanisms including alkylating agents, antimetabolites, antimitotics, and monoclonal antibodies (43). However, as far as we know, no one has attempted to treat cancer through modulation of protein assembly, giving our work with PAV-617 and PAV-951 the potential to be both risky and rewarding. While the animal toxicity of PAV-617 and PAV-951 is higher than would be ideal for the clinic, their anti-tumor activity is already on par with existing cancer drugs and a handful of FDA approved cancer drugs have comparable toxicity gauged by mouse MTD—cisplatin has a MTD of 6 mg/kg, Doxorubicin has a MTD of 10 mg/kg, and Vinorelbine has a MTD of 10 mg/kg- and many other cancer drugs are administered to patients despite adverse effects because of the urgency of their condition (44,45). Since PAV-617 and PAV-951 are early compounds, further optimization will likely yield chemical analogs with substantially reduced toxicity and further increased activity. Indeed, we have already identified chemical analogs that demonstrate substantially increased safety in mice without losing anti-cancer activity in cells (see **Supplemental Figure 3**). We hypothesize that driving SAR toward compounds that are selective for restoring feedback loops of protein homeostasis will continue to improve the therapeutic indexes because the mechanism-of-action does not inherently pose a risk of toxicity/collateral damage towards normal cells.

Molecular genetic tools such as CRISPR, siRNA knock down, and even use of recombinant protein for protein-protein interaction studies, are unable to parse out the post-translational heterogeneity introduced into proteins as part of normal and aberrant biochemical pathways. The methods applied here are able to do so, as evidenced by the small fraction of the total of specific proteins such as KAP-1 found in the target multi-protein complex. It is perhaps not surprising that new tools, and the new targets they allow to be detected, make possible a path to drugs with novel properties. Further work will clarify whether, as we hypothesize, the transience of our targets reflects their involvement as a molecular basis for homeostasis. This conclusion, supported by the consequences of protein assembly modulator treatment both here for cancer and previously for viruses, frames future experiments to better understand this novel approach to more physiological disease therapeutics.

## Supplemental Figures

Synthesis of A. PAV-617 and B. PAV-617 resin

**Supplemental Figure 1.**
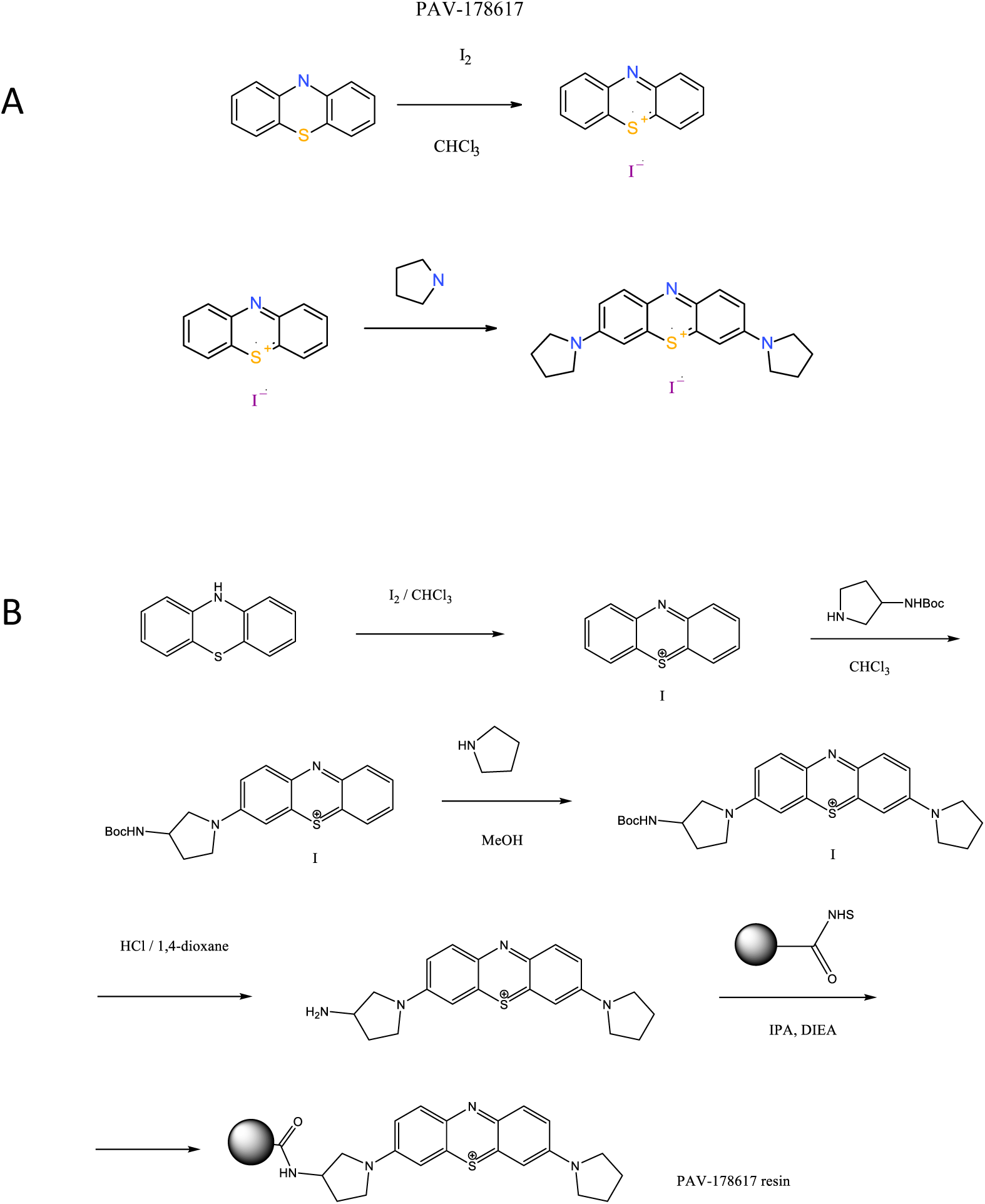
Structure and synthetic scheme of PAV-617 and its resin. **Supplemental Figure 1A** shows the chemical synthesis of PAV-617. **Supplemental Figure 1B** shows the chemical synthesis of PAV-617 resin.

**Supplemental Figure 2.**
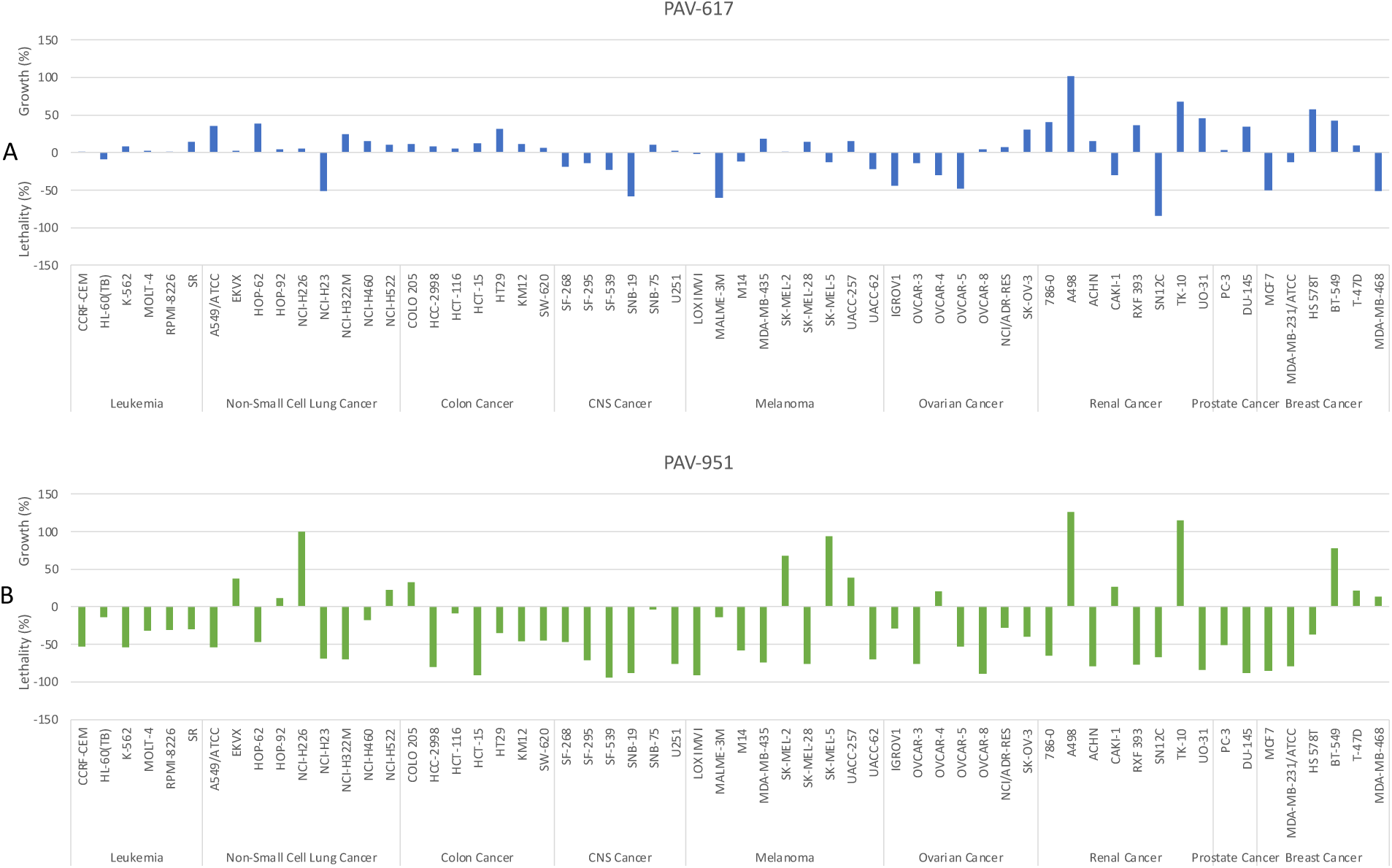
NCI-60 Cell screen with (A) PAV-617 and (B) PAV-951. Growth percent of each cell line after 48 hr treatment with 2.5 uM PAV-617 or 2.5 uM PAV-951 is relative to the vehicle treated control and the number of cells at time zero. Values between 0 and 100 represent percent growth inhibition and values between 0 and -100 represent percent cellular lethality. Figure A shows PAV-617 exhibiting greater than 50% growth inhibition on 37 cell lines and cellular lethality on 20 cell lines. Figure B shows PAV-951 exhibiting greater than 50% growth inhibition on 9 cell lines and cellular lethality on 45 cell lines. These findings corroborate and extend the initial findings.

**Supplemental Figure 3.**
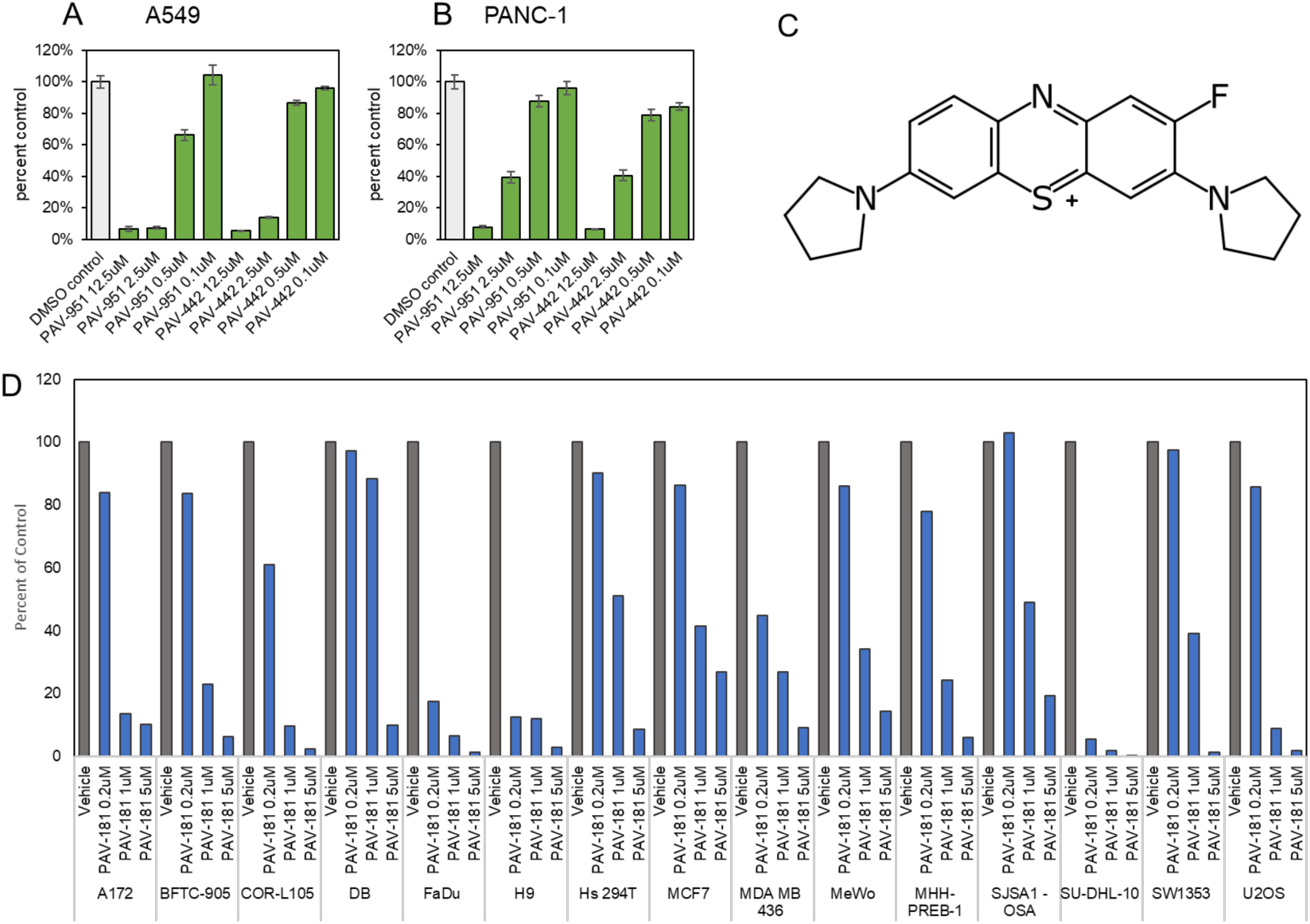
SAR optimization of the PAV-951 series. **Supplemental Figures 2A** and **2B** show comparable anticancer activity of PAV-951 and its less toxic (6x higher mouse MTD) chemical analog, PAV-442, on reducing growth of A549 lung cancer and PANC-1 pancreatic cancer tumor lines relative to a vehicle-only control. Supplemental Figure 2C shows the disclosed chemical structure of PAV-181, a chemical analog of PAV-617. Supplemental Figure 2D shows the activity of PAV-181 in at 5uM, 1uM, and 0.2uM against A172 (male human glioma), BFTC-905 (female human urinary bladder transitional cell carcinoma), COR-L105 (male human lung adenocarcinoma), DB (male human b-cell lymphoma), FaDu (male human pharynx squamous cell carcinoma), H9 (male human t-cell lymphoma), Hs 294T (male human melanoma), MCF7 (female human breast cancer), MDA MB 436 (female human breast cancer), MeWo (male human melanoma), MHH-PREB-1 (male human b-cell lymphoma), SJSA1-OSA (male human osteosarcoma), SU-DHL-10 (male human b-cell lymphoma), SW1353 (female human chondrosarcoma), and U-2 OS (female human osteosarcoma).

**Supplemental Figure 4.**
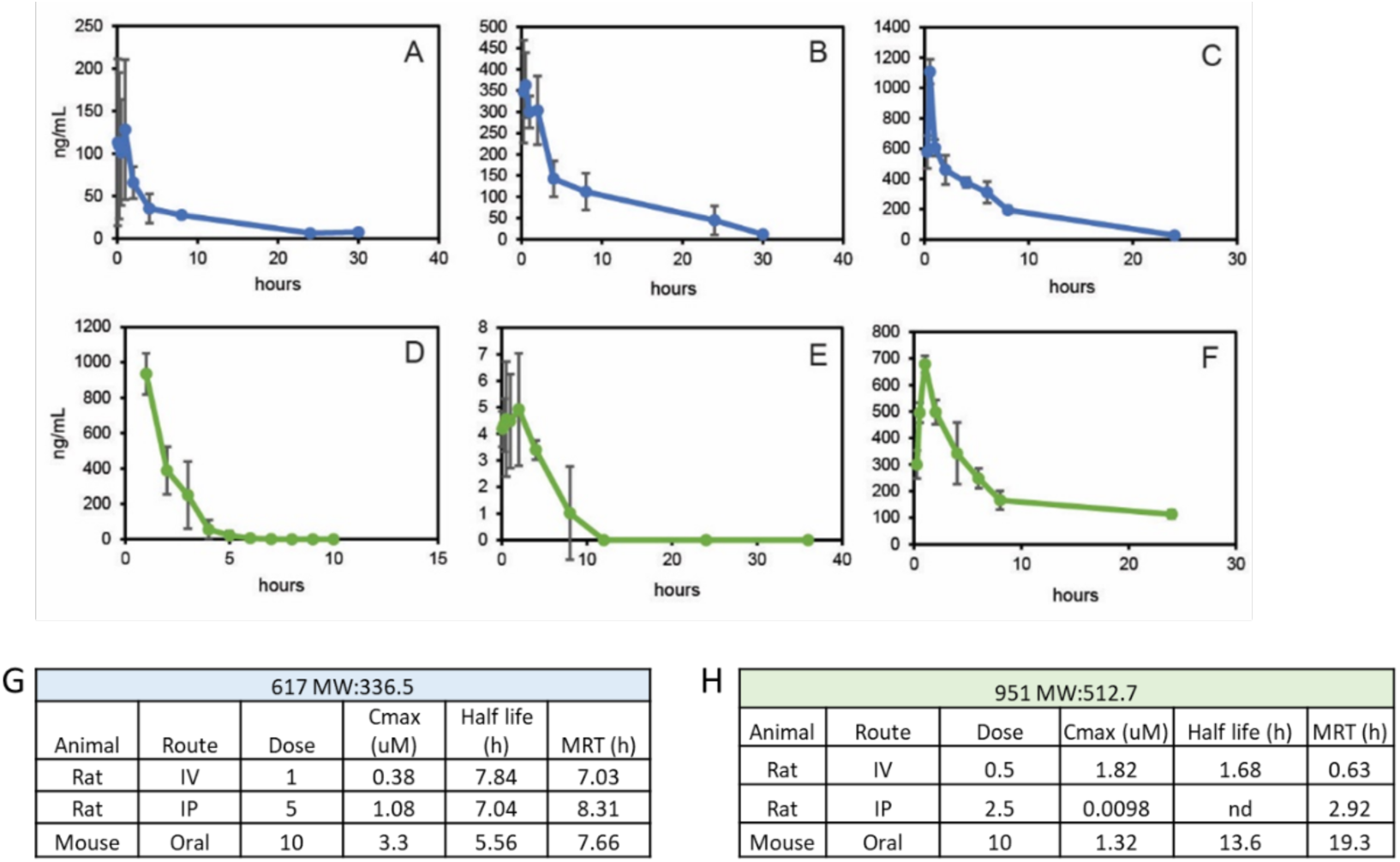
Summary of PAV-617 and PAV-951 PK properties. **Supplemental Figures 3A-F** show plasma concentration over time for PAV-617 and PAV-951 when given to animals via three different routes of administration. Randomized treatment groups of four male Sprague Dawley rats or three CD1 mice were administered vehicle, PAV-617 (panel A-C), or PAV-951 (panel D-F) either IV (A, D), IP (B, E), or orally (C, F) and blood samples were collected before injection as well as at different time points after dosing. The concentration of compound in the plasma at different time points was measured by LC MS/MS. **Supplemental Figure 4G** summarizes PK properties observed for PAV-617 based on animal, administration route, and dose. **Supplemental Figure 4H** summarizes the PK properties observed for PAV-951 based on animal, administration route, and dose. The maximum concentration (Cmax) was determined to be the highest measured concentration within a dataset. The half-life and mean residence time (MRT) were calculated from the values determined over time.

**Supplemental Figure 5.**
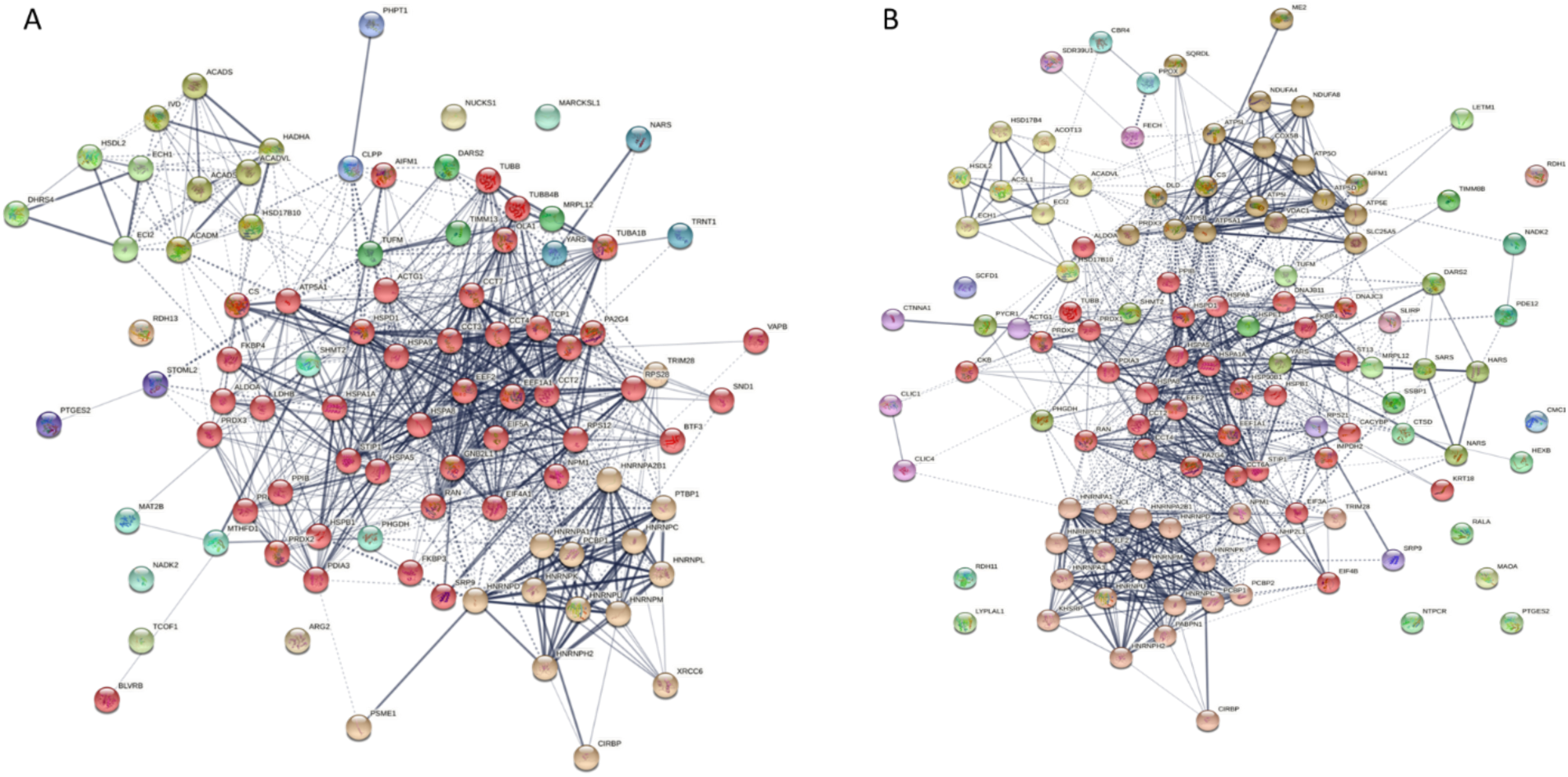
Known protein-protein interactions among PAV-617 and PAV-951 eluate components. **Supplemental Figures 5A** and **5B** show string-diagram analyses of the protein-protein interaction network of proteins identified in **Figures 5A-C** by tandem MSMS as comprising the PAV-617 (**Supplemental Figure 5A**) and PAV-951 (**Supplemental Figure 5B**) targets. Proteins were entered into the string database (string-db.org). The confidence mode view is shown where the confidence level was set to 0.4 (medium) and MCL clustering was applied with the inflation parameter of 3. The dotted lines represent edges between different clusters. The thickness of the lines represents the degree of confidence for each protein-protein interaction.

**Supplemental Figure 6.**
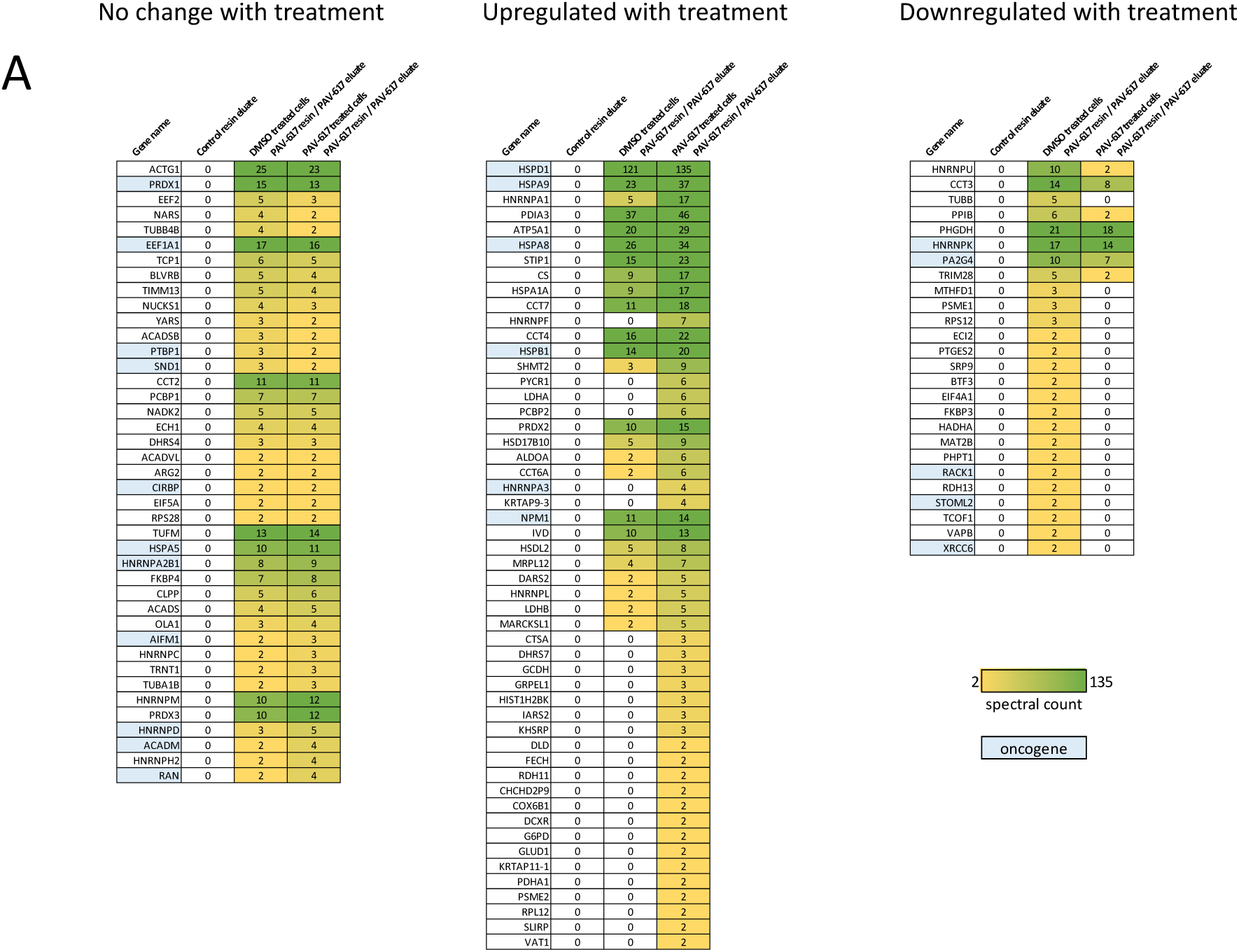
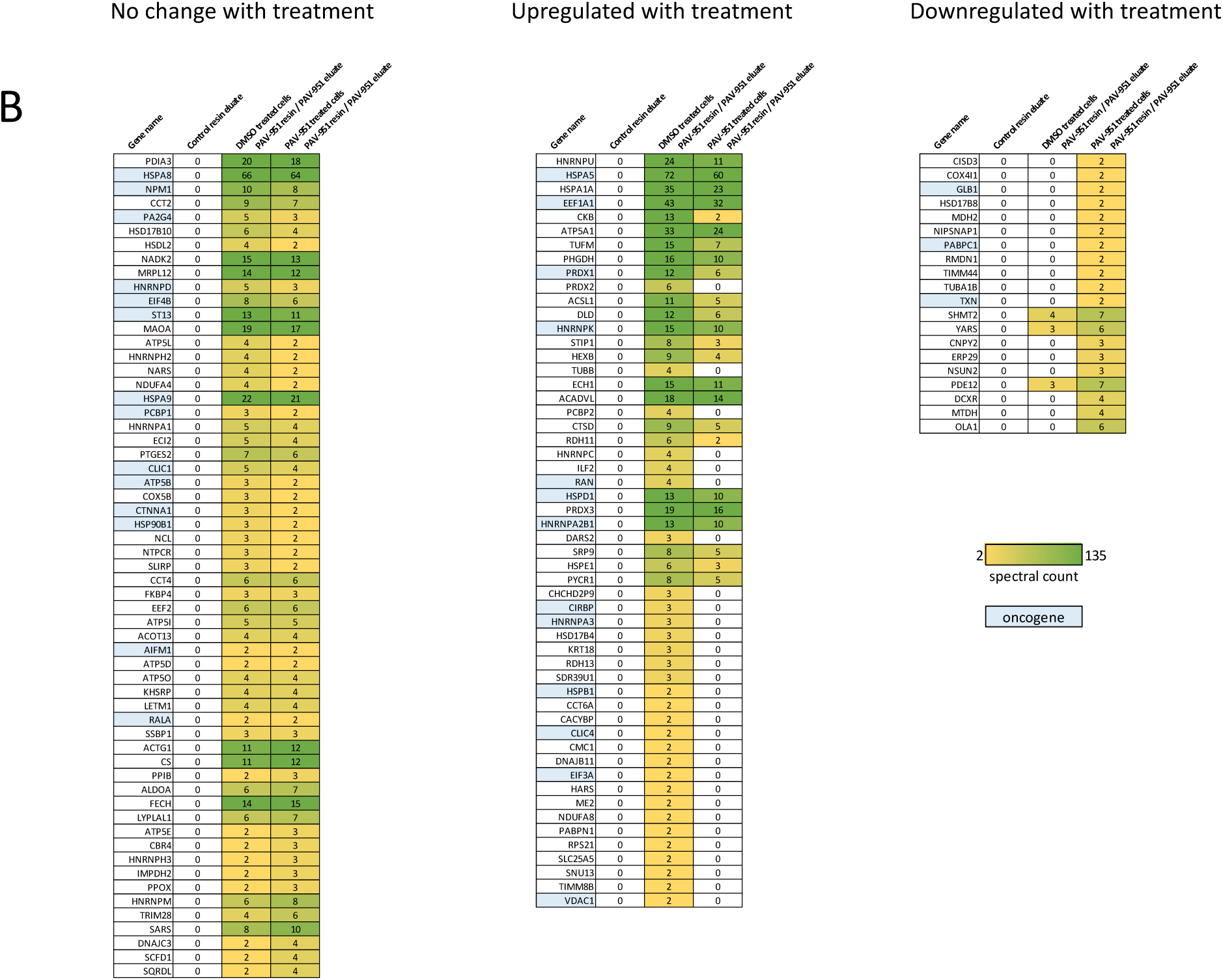
The same data analyzed in terms of selective vs shared proteins between the drug resin in Figure 5 is here sorted by upregulation, downregulation or no change, with drug treatment for PAV-617 (A) and PAV-951 (B). The oncogenes and other cancer-related proteins are highlighted here in blue.

**Supplemental Figure 7.**
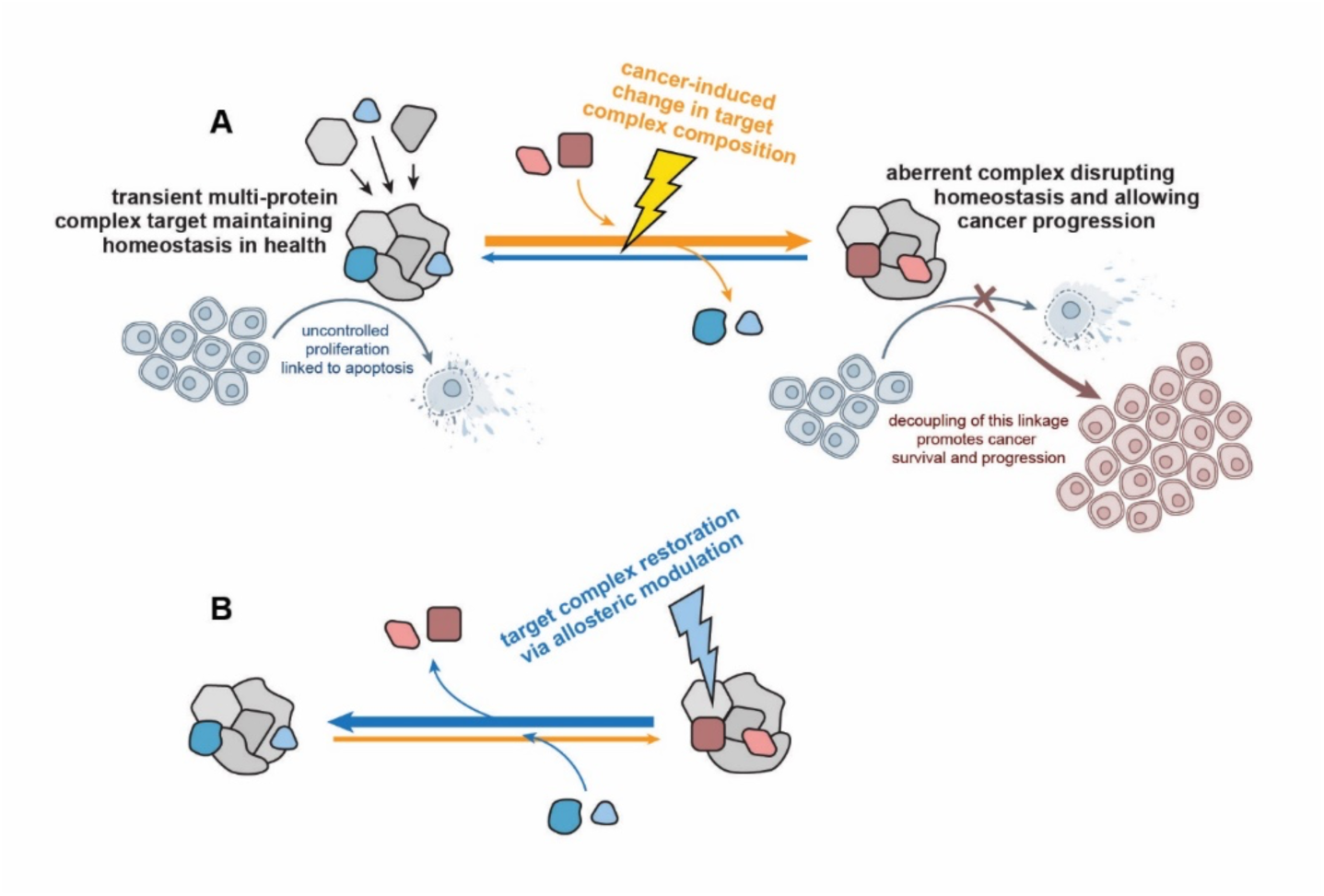
Cartoon diagram of proposed mechanism of action of assembly modulating compounds. **Supplemental Fig. 7A** The proposed model where a normal multi-protein complex that plays a role mediating the linkage between uncontrolled proliferation and apoptosis is modified into an aberrant multi-protein complex at an early, precancerous stage allowing cancer progression rather than homeostatic elimination. **Supplemental Figure 7B** shows the proposed mechanism where treatment with a protein assembly modulating compound restores the original multi-protein complex and its homeostatic functions, including elimination of the cancer. Allosteric modulation is indicated as a means by which these changes may be induced(46).

## Materials and Methods

### Lead Contact and Materials Availability

Further information and requests for resources and reagents should be directed to and will be fulfilled by the Lead Contact Vishwanath R. Lingappa.

Use of unique compounds PAV-617 and PAV-951 and their stable derivatives may be available upon request by the Lead Contact if sought for experimental purposes under a valid completed Materials Transfer Agreement.

The number of replicates carried out for each experiment is described in the figure/table legends.

### Chemical Synthesis

All compounds synthesized were confirmed by LCMS with purity typically > 98%

**Synthesis of PAV-617** (see **Supplemental Fig. 4A**)

#### Phenothiazin-5-ium tetraiodide hydrate

Phenothiazin-5-ium tetraiodide hydrate: a solution of phenothiazine (4.98 g, 25 mmol) in anhydrous chloroform (50 ml) was stirred at 50C and the solution of iodine (12.7 g, 50 mmol) in CHCl3 (250 ml) was added dropwise over 4h. The resulting dark solution was stirred for an additional 3h at 50C, monitored by TLC. After the disappearance of the starting material, the resulting precipitate was filtered, washed with a copious amount of chloroform, dried overnight in vacuo to afford a dark solid (13.9 g, 74%).

#### 3,7-Di(pyrrolidin-1-yl)phenothiazinium iodide

A solution of phenothiazin-5-ium tetraiodide hydrate (2.8 g, 3.6 mmol) in mixture acetonitrile/methanol (50 ml) and pyrrolidine (710 mg, 10 mmol) was stirred for 4 h at room temperature. The resulting mixture was concentrated to dryness and purified by flash chromatography using the methanol-chloroform gradient to purify the desired compound.

### Synthesis of PAV-617 resin and crosslinker **(see** Supplemental Fig. 4B)

#### To 6-(tert-butoxycarbonylamino)-2-(9H-fluoren-9-ylmethoxycarbonylamino)hexanoic acid

[468mg (1mmol)] was added 3ml of 4N HCl and the resulting solution was stirred for 15min at room temperature. The mixture was rotary evaporated to a residue which was then sequentially treated with DMF (4ml), Diazirine 10 [128mg (1mmol)], DIEA [347ul (2mmol)] and HATU [380mg (1mmol)]. After stirring for 30min at room temperature the mixture was diluted with EtOAc (20ml) and washed with water (3x). The organic layer was dried (Mg2SO4) and the solvent removed affording the Diazirine acid 11 which was used as is.

To the Diazirine acid 11 [24mg (0.05mmol)] was added tert-Butyl 4-aminobutanoate [8mg (0.05mmol)], DMF (1ml), DIEA [35ul (0.2mmol)] and HATU [19mg (0.05mmol)]. After stirring for 30min at room temperature the mixture was diluted with EtOAc (5ml) and washed with water (3×1ml). The organic layer was dried (Mg2SO4) and the solvent removed affording the Diazirine amide XX which was used as is.

The crude Diazirine amide XX was next treated with a 50% Diethylamine / DMF solution (1ml) for 1h at room temperature with mild agitation. The solvent and Diethylamine were then removed using high vacuum overnight. The residue was the treated with DMF (1ml), DIEA (one drop) and Biotin-PEG2-NHS [25mg (0.05mmol)]. The mix was stirred for 12 at room temperature and again the solvent was removed using high vacuum affording the crude Biotinylated Diazirine YY.

The crude Biotinylated Diazirine YY was next treated with 4N HCl in Dioxane (1ml) for 30min at room temperature then the mixture was rotary evaporated to dryness. The residue was then dissolved in DMF (1ml) and then treated with DIEA [35ul (0.05mmol)], the Phenothiazinium chloride salt ZZ [21mg (0.05mmol)] and HATU [19mg (0.05mmol)]. After 30min the solvent was removed using high vacuum. The residue was the purified via reverse phase chromatography (0.1% TFA in ACN / 0.1% TFA in water) which after lyophilization afforded 24mg (47% overall yield) of the desired PAV-617 crosslinker (XYZ).

### Experimental Models and Subject Details

#### Animal models

Maximum tolerated dose (MTD) studies were conducted using female Balb/C mice, aged 8-10 weeks or female CD1 mice, aged 5-6 weeks. Treatment groups were made up of 3 animals each, unless otherwise noted, and dosing regimens for disclosed data is provided. Animals were sacrificed at the end of the study period using an overdose of CO2. MTD studies were conducted at Vipragen Biosciences Private Limited or Radiant Research Services Private Limited in accordance with the Committee for the Purpose of Control and Supervision of Experiments on Animals (CPCSEA) guidelines and Animal Research: Reporting of In Vivo Experiments (ARRIVE) guidelines.

Pharmacokinetic (PK) studies were conducted in male Sprague Dawley Rats aged 8-10 weeks or male CD1 mice aged 5-6 weeks. Treatment groups were made up of 4 animals each and dosing regimens for disclosed data is provided. Animals were sacrificed at the end of the study period using an overdose of CO2. PK studies were conducted at Vipragen Biosciences Private Limited in accordance with the Committee for the Purpose of Control and Supervision of Experiments on Animals (CPCSEA) guidelines and Animal Research: Reporting of In Vivo Experiments (ARRIVE) guidelines.

Tumor xenograft studies were conducted using female Athymic Nude mice, strain CrTac: Ncr-Foxn1^nu^, aged 6-8 weeks. Tumor transplantation occurred through subcutaneous injection of a 0.1mL cell suspension containing 1 to 5×10^6^ A549 lung cancer cells obtained from ATCC in Matrigel in PBS into the left flank region of the mice. Treatment groups were made up of 6 animals each and dosing regimens for disclosed data is provided. Animals were sacrificed at the end of the study period using an overdose of isoflurane anesthesia. Both xenograft studies were conducted at Anthem Biosciences Private Limited in Bangalore, India and were approved by the Institutional Animal Ethics committee (IAEC) of Anthem Biosciences in accordance with the Committee for the Purpose of Control and Supervision of Experiments on Animals (CPCSEA) guidelines and Animal Research: Reporting of *in vivo* Experiments (ARRIVE) guidelines.

#### Cell lines

The *in vivo* xenograft study utilized A549 (male human lung cancer) and HT-29 (female human colon cancer) cell lines. Cells were grown under sterile conditions. These studies were conducted at Anthem Biosciences Private Limited in Bangalore, India.

The Human tumor cell proliferation assay used A172 (male human glioma), BFTC-905 (female human urinary bladder transitional cell carcinoma), COR-L105 (male human lung adenocarcinoma), DB (male human b-cell lymphoma), FaDu (male human pharynx squamous cell carcinoma), H9 (male human t-cell lymphoma), Hs 294T (male human melanoma), MCF7 (female human breast cancer), MDA MB 436 (female human breast cancer), MeWo (male human melanoma), MHH-PREB-1 (male human b-cell lymphoma), SJSA1-OSA (male human osteosarcoma), SU-DHL-10 (male human b-cell lymphoma), SW1353 (female human chondrosarcoma), and U-2 OS (female human osteosarcoma) cell lines. This study was conducted by Eurofins Scientific as part of their OncoPanel^TM^.

The monkey pox virus infectious virus assay used BSC-40 cells (nonhuman primate kidney) and MPXV Zaire 79 strain. These studies were conducted by the United States Army Medical Research Institute of Infectious Diseases on Fort Detrick, Maryland.

Human immunodeficiency virus infectious virus assay used MT-2 cells (female human t-cell leukemia) and the NL4-3 Rluc reporter virus. These studies were conducted at the University of Washington in Seattle, Washington.

The *in vitro* screens for apoptosis, high-density/ low-density activity, and cell growth recovery utilized the LNCaP C-33 (male human prostate cancer), LNCaP C-81 (male human prostate cancer), CHO-K1 (Chinese Hamster ovary), and Hennes 20 (Chinese Hamster ovary) cell lines. Cells were grown under sterile conditions. These studies were conducted at Prosetta Biosciences in San Francisco, California.

The *in vitro* drug resin affinity chromatography and photocrosslinking experiments utilized A549 (male human lung cancer) or LNCaP C-33 (male human prostate cancer) cell line. Cells were grown under sterile conditions. Sterile conditions were not maintained once cells were harvested for *in vitro* experiments. These studies were conducted at Prosetta Biosciences in San Francisco, California.

#### National Cancer Institute Sixty Cell Line Screen

The human tumor cell lines of the cancer screening panel are grown in RPMI 1640 medium containing 5% fetal bovine serum and 2 mM L-glutamine. For a typical screening experiment, cells are inoculated into 96 well microtiter plates in 100 μL at plating densities ranging from 5,000 to 40,000 cells/well depending on the doubling time of individual cell lines. After cell inoculation, the microtiter plates are incubated at 37° C, 5 % CO2, 95 % air and 100 % relative humidity for 24 h prior to addition of experimental drugs.

After 24 h, two plates of each cell line are fixed *in situ* with TCA, to represent a measurement of the cell population for each cell line at the time of drug addition (Tz). Experimental drugs are solubilized in dimethyl sulfoxide at 400-fold the desired final maximum test concentration and stored frozen prior to use. At the time of drug addition, an aliquot of frozen concentrate is thawed and diluted to twice the desired final maximum test concentration with complete medium containing 50 μg/ml gentamicin

Following drug addition, the plates are incubated for an additional 48 h at 37°C, 5 % CO2, 95 % air, and 100 % relative humidity. For adherent cells, the assay is terminated by the addition of cold TCA. Cells are fixed *in situ* by the gentle addition of 50 μl of cold 50 % (w/v) TCA (final concentration, 10 % TCA) and incubated for 60 minutes at 4°C. The supernatant is discarded, and the plates are washed five times with tap water and air dried. Sulforhodamine B (SRB) solution (100 μl) at 0.4 % (w/v) in 1 % acetic acid is added to each well, and plates are incubated for 10 minutes at room temperature. After staining, unbound dye is removed by washing five times with 1 % acetic acid and the plates are air dried. Bound stain is subsequently solubilized with 10 mM trizma base, and the absorbance is read on an automated plate reader at a wavelength of 515 nm. For suspension cells, the methodology is the same except that the assay is terminated by fixing settled cells at the bottom of the wells by gently adding 50 μl of 80 % TCA (final concentration, 16 % TCA). Using the seven absorbance measurements [time zero, (Tz), control growth, (C), and test growth in the presence of drug at the five concentration levels (Ti)], the percentage growth is calculated at each of the drug concentrations levels. Percentage growth is calculated as:

[(Ti-Tz)/(C-Tz)] × 100 for concentrations for which Ti>/=Tz
[(Ti-Tz)/Tz] × 100 for concentrations for which Ti<Tz.

#### *In vitro* experiments

Drug resin affinity chromatography and photocrosslinking experiments, and SDS-PAGE/ silver stain/Western Blot analysis of the results, were conducted by Prosetta Biosciences in San Francisco, California under conditions described in figure legends. Results from disclosed *in vitro* experiments were repeated in triplicate unless otherwise stated. Mass spectrometry analysis of samples were conducted by MS Bioworks in Ann Arbor, Michigan.

## Method and Analysis Details

### Monkey pox infectious virus assay

BSC-40 cells of 95% confluence in 24-well plates were infected with100 pfu of MPXV Zaire 79 diluted in Eagle’s Minimum Essential Medium with 2% fetal bovine serum and incubated in 37 degrees Celsius in 5% CO_2_ for 1 hour. The viral inocula were removed and replaced with the test compounds in six half log dilutions (0.1 ml per well) and the cells were overlaid with 1% methylcellulose in growth media (1 ml per well). The media and virus control cells received growth medium containing 1% methylcellulose. After three days of infection, when plaques appeared, cells were stained with crystal violet for an hour and then washed with water and dried overnight. The plaques were counted the next day and virus-only wells were compared with the compound-added wells to determine percentage protection. Infected cells were stained with crystal violet and viral plaques were counted. Averages and standard deviation for plaques observed under different treatment conditions were calculated in Microsoft Excel and graphed as the percent inhibition in PAV-617 treated cells compared to untreated cells.

### Human immunodeficiency virus infectious virus assay

MT-2 cells were preseeded in 96-well plates in 100 ul of complete RPMI. Multiple concentrations of PAV-951 were serially diluted in DMSO then into an infection media prepared by diluting NL4-3 Rluc virus stock to 400 IU/100 ul with complete RPMI, which was transferred onto the MT-2 cells with a final MOI of 0.02 and final DMSO concentration of 1% in infected places. One well received DMSO only, instead of PAV-951, and one well received medium only for normalization and background collection. Cells were incubated at 37 degrees Celsius for 96 hours. 100ul of medium was removed and discarded and 10 ul of 15 uM EnduRen luciferase substrate was added to each well, followed by incubation for 1.5 hours at 37 degrees Celsius. Plates were read on a luminescence plate reader. Bioluminescence intensity was read on a Synergy H1 BioTek plate reader. Averages and standard deviation for viral titer observed under different treatment conditions were calculated in Microsoft Excel and graphed as the percent inhibition in PAV-951 treated cells compared to untreated cells.

### Apoptosis Screen

A 96 well plate was seeded with Hennes 20 cells at 500 cells per well, CHO-K1 cells at 500 cells per well, LNCaP C-33 cells at 2000 cells per well, and LNCaP C-81 cells at 2000 cells per well. Cells were grown in 100uL minimum essential media for three days then three wells of each cell line received treatment with 1% DMSO. 12 hours after drug treatment, a mixture of 25 ul media and 25 ul Apo-ONE reagent (Promega) was added then the plate was covered and placed on a shaker at room temperature for six hours. The plate was read on a microplate reader for fluorescence at 499/521. Values were averaged and standard deviations were calculated for each triplicate condition and graphed on Microsoft Excel.

### High density/ low density assay

Two 96 well plates were seeded with Hennes 20 cells in parallel where one was plated at a density of 500 cells/well and the other was plated at a density of 15,000 cells/well. 90 ul of minimum essential media was added to each well and plates were placed in a 37 degrees Celsius incubator for 24 hours. The next day, 10ul of media containing dilutions of compound in DMSO were added to each plate in triplicate with final concentrations of 0.025 uM PAV-617, 0.05 uM PAV-617, 0.1 uM PAV-617, 0.5 uM PAV-617, 0.02 uM PAV-951, 0.3 uM PAV-951, 0.4 uM PAV-951, or 0.5 uM PAV-951. Six wells on each plate received 10ul of media containing only DMSO. Each well was gently mixed 5 times with a 100ul pipette. Plates were incubated at 37 degrees Celsius for 72 hours then 10 uL of alamarBlue was added to each well. Wells were mixed 5 times then incubated at 37 degrees Celsius for 72 hours. Plates were then read at 530/590. Values were averaged and standard deviations were calculated for each triplicate condition and graphed on Microsoft Excel.

### Cell growth recovery assay

A 96 well plate was seeded with either Hennes 20 or LNCaP C-33 cells at 500 cells/well in 90 uL of minimum essential media and incubated at 37 degrees Celsius for 24 hours. 0.5% DMSO was diluted in media and added to 6 control wells for each plate. PAV-617 was diluted in media and added to three wells at a concentration of 0.3 uM. PAV-951 was diluted in media and added to concentration of 0.4 uM. After 24 hours of PAV-617 treatment or 6 hours of PAV-951 treatment, the medium containing compound was removed and replaced with fresh media. After 72 hours (day 5), plates were assayed with alamarBlue and fluorescence was read at 530/590. The medium containing alamarBlue was removed and replaced with fresh media. After another 72 hours (day 8) plates were assayed with alamarBlue and fluorescence again, then medium containing alamarBlue was removed and replaced with fresh media. After a final 72 hour incubation (day 11) plates were assayed with alamarBlue one more time. Average fluorescence for each day and treatment condition were plotted on Microsoft Excel with standard deviation calculated to provide error bars.

### Human tumor cell proliferation assay

A panel of human tumor cell lines (A172, BFTC-905, COR-L105, DB, FaDu, H9, Hs 294T, MCF7, MDA MB 436, MeWo, MHH-PREB-1, SJSA1-OSA, SW1353, and U2OS) were grown in RPMI 1640, 10% FBS, 2 mM L-alanyl-L-glutamine, 1 mM Na pyruvate. Cells were seeded into 384-well plates and incubated in a humidified atmosphere with 5% CO2 at 37C. After 24 hours of incubation DMSO, PAV-617, or PAV-951 was added at concentrations of 5 uM, 1 uM, and 0.2 uM and plates were incubated for 3 days. Then cells were lysed with CellTiter-Glo (Promega) which generates a bioluminescence signal relative to ATP levels and is used as a measurement of viable cells. Bioluminescence was read by a PerkinElmer Envision microplate reader. Bioluminescence intensity was measured by a PerkinElmer Envision microplate reader and transformed to a percent of control (POC) using the formula: POC=(Ix/I0)*100, where Ix is the whole well signal intensity at a given treatment, and I0 is the average intensity of the untreated vehicle wells. Values were averaged for each triplicate condition and graphed on Microsoft Excel.

### Screening for SAR

A549 and PANC1 cells were seeded at a density of 1,250 cell/well in 100ulo of media/well. Plates were incubated at 37 degrees Celsius and 5% CO2 for 24 hours. DMSO, PAV-951, or its analog PAV-442 were added to media at final concentrations of 0.1 uM, 0.5uM, 2.5 uM, or 12.5uM in 0.5% DMSO. The treated plates were incubated at 37 degrees celsius for 72 hours than analyzed by AlamarBlue.

### Mouse maximum tolerated dose determination

For the intraperitoneal MTD study, female Balb/c mice aged 8-10 weeks were randomly divided into treatment groups with three animals per group. Animals in each treatment group were weighed and received one IP injection of 0.1-0.15mL containing either vehicle (10% DMSO, 45% propylene glycol, 45% sterile water), 1mg/kg PAV-617, 2 mg/kg PAV-617, 5 mg/kg PAV-617, 10 mg/kg PAV-617, 1mg/kg PAV-951, 2.5 mg/kg PAV-951, 5mg/kg PAV-951 or 10 mg/kg PAV-951. Animals were observed from day 0 until day 3 for clinical signs of toxicity. Animals were euthanized after 72 hours and were examined externally and internally by a pathologist for abnormalities in organ weight and tissue damage. Blood samples were sent for a complete blood count bioanalysis. MTD was determined to be the dose at which no signs of toxicity were observed by any parameters.

For the oral MTD study, female CD1 mice aged 5-6 weeks were given either an oral dose of vehicle (10% DMSO, 45% propylene glycol, 45% sterile water) or either 10 mg/kg or 20 mg/kg of PAV-617 or PAV-951. The vehicle and 20 mg/kg groups had three animals each, while the 10 mg/kg groups only had one animal. Animals were observed for clinical signs and after a week they were euthanized and examined externally and internally by a pathologist for changes related to toxicity.

### Pharmacokinetics studies

For the IP and IV PK studies, male Sprague Dawley rats aged 8-10 weeks were randomly divided into treatment groups with four animals per group. Animals in each treatment group were weighed and received one 2.4 mL intravenous dose of either vehicle (10% DMSO, 45% propylene glycol, 45% sterile water), 1mg/kg PAV-617, or 0.5 mg/kg PAV-951, or one intraperitoneal dose of either vehicle (100% labrasol), 5mg/kg PAV-617 or 2.5 mg/kg PAV-951 . Blood was collected from a pre-cannulated line before dosing, and subsequently 15 minutes, 30 minutes, 1 hour, 2 hours, 4 hours, 8 hours, 12 hours, 24 hours, and 30 hours post-dosing. Concentration of drug in the plasma over time was measured using a Waters Acquity TQD LCMS/MS. Maximum concentration (Cmax) was determined to be the maximum concentration detected in a dataset. Half life, area under the curve, and mean residence time were calculated with Phoenix WinNolin software.

For the oral PK studies, male CD1 mice aged 5-6 weeks were randomly divided into treatment groups with three animals per group. Animals in each treatment group were weighed and received, via oral gavage needle, either one oral dose of vehicle (10% DMSO, 45% propylene glycol, 45% sterile water), 10 mg/kg PAV-617, or 10 mg/kg PAV-951. Blood was collected from a pre-cannulated line before dosing, and subsequently 2 minutes, 5 minutes, 15 minutes, 30 minutes, 1 hour, 2 hours, 4 hours, 6 hours, 8 hours, and 24 hours post-dosing. Concentration of drug in the plasma over time was measured using a Waters Acquity TQD LCMS/MS. Maximum concentration (Cmax) was determined to be the maximum concentration detected in a dataset. Half life, area under the curve, and mean residence time were calculated with Phoenix WinNolin software.

### A549 xenograft studies

A549 cells growing in RPMI-1640 medium were suspended with Matrigel in PBS. 0.1 ml of cell suspension containing 1 × 10^6^ cells were injected subcutaneously into the left flank region of female, 6-8 weeks old nude mice (CrTac: Ncr-Foxn1nu). After 30 days of tumor establishment, mice were divided randomly into treatment groups. In the PAV-617 study, 6 animals were treated with vehicle only (10% DMSO, 10% propylene glycol, 80% sterile water) by IP once daily, 6 animals were treated with 100 mg/kg Gemcitabine Hydrochloride by IP twice weekly, and 6 animals were treated with 10 mg/kg PAV-617 by IP once daily for 28 days. In the PAV-951 study, 6 animals were treated with vehicle only (10% DMSO, 10% propylene glycol, 80% sterile water) by IV once daily, 6 animals were treated with 100 mg/kg Gemcitabine by IV twice weekly, and 6 animals were treated with 1.5 mg/kg PAV-951 by IV once daily for 14 days. In both studies, mice were weighed and their tumors were measured using a digital Vernier caliper. Tumor volume was calculated using the formula: (L × W^2^)/2 where L is the largest diameter and W is the smallest diameter of the tumor. Statistical analysis was performed using Graph Pad Prism (Ver. 5.03). Statistical analysis of tumor growth inhibition between the Control and Treated groups was performed by using One-way ANOVA followed by Dunnett’s test.

### HT-29 xenograft study

HT-29 cells growing in HBSS medium were suspended in Matrigel. 0.1 mL of the cell suspension containing 5×10^6^ cells were injected subcutaneously into the left flank region of male, 6-8 week old SCID mice. After tumor establishment, animals were divided randomly into groups of 6 and treated daily by IV with vehicle daily for 17 days, 3 mg/kg PAV-951 three times per week for 17 days, or treated by IP with 60 mg/kg Irinotecan every 4 days for 14 days. Tumor volume was calculated using the formula: (L × W^2^)/2 where L is the largest diameter and W is the smallest diameter of the tumor. Statistical analysis was performed using Graph Pad Prism (Ver. 5.03). Statistical analysis of tumor growth inhibition between the Control and Treated groups was performed by using One-way ANOVA followed by Dunnett’s test.

### Drug Resin affinity chromatography

LNCaP C-33 cells were grown in minimum essential media (UCSF) with 10% FBS and 1% Penstrep for 24 hours then treated with 500nM PAV-617, 500nM PAV-951, or DMSO for 22 hours. Cells were scraped into cold phosphate buffered saline (PBS) (10mM sodium phosphate, 150 mM sodium chloride pH 7.4), then spun at 1,000 rpm for 10 minutes until pelleted. The PBS was decanted and the pellet resuspended in a low salt buffer (10mM HEPES pH 7.6, 10mM NaCl, 1mM MgAc with 0.35% Tritonx100) then centrifuged at 10,000 rpm for 10 minutes at 4o C. The post-mitochondrial supernatant was removed and adjusted to a concentration of approximately 10 mg/ml and equilibrated in a physiologic column buffer (50 mM Hepes ph 7.6, 100 mM KAc, 6 mM MgAc, 1 mM EDTA, 4mM TGA). In some conditions, the extract was supplemented with an energy cocktail (to a final concentration of 1mM rATP, 1mM rGTP, 1mM rCTP, 1mM rUTP, and 5 ug/mL creatine kinase). 30 ul or 230 ul of extract was then incubated for one hour at either 4°C or 22°C on 30 ul or 230 ul of affigel resin coupled to either PAV-617, PAV-951, or a 4% agarose matrix (control). The input material was collected and the resin was then washed with 3 ml column bufffer. The resins were eluted overnight at either 4 °C or at 22°C in 100ul or 330ul column buffer containing either 100uM PAV-617 or 100uM PAV-951 or DMSO, with or without the energy cocktail. Eluates were run on SDS-PAGE with samples for silver stain and/or western blot or sent for mass spectrometry analysis.

### Chemical photocrosslinking

A549 extract was prepared as above then adjusted to a protein concentration of approximately 1 mg/ml in column buffer containing 0.01% triton and supplemented with the energy cocktail (to a final concentration of 1mM rATP, 1mM rGTP, 1mM rCTP, 1mM UTP, and 5 ug/mL creatine kinase). Photocrosslinker analogs of PAV-617 and PAV-951 chemically modified to contain biotin and a diazirine group or 1% DMSO were added to 67 ul of extract at 100uM, incubated for one hour at 22°C followed by 20 minutes on ice, then exposed to ultraviolet light for 5 minutes at 22°C. After crosslinking, samples were divided in two 30 ul aliquots and one set was denatured by adding 5 ul of 10% SDS, 0.625 ul DTT, and boiling for 5 minutes. Both native and denatured aliquots were then diluted in 800 ul column buffer containing 0.1% triton. 2.5 ul of magnetic streptavidin beads (Pierce) were added to all samples and mixed for one hour at room temperature to capture all biotinylated proteins and co-associated proteins. Samples were placed on a magnetic rack to hold the beads in placed and washed three times with 800 ul of column buffer containing 0.1% triton. After washing, beads were resuspended in 80 ul of gel loading buffer containing SDS and analyzed by western blot or blot for affinity purified streptavidin. Samples were analyzed by western blot.

### Silver Stain

SDS/PAGE gels were incubated overnight in a fixative (50% methanol, 10% acetic acid, 40% water), then for an hour in 50% methanol (done as two washes), and an hour in water (done as two washes). The gels were sensitized in 0.02% sodium thiosulfate for one minute then washed twice for 30 seconds with water. The gels were incubated for 30 minutes in cold 0.1% silver nitrate with 0.02% formaldehyde then washed twice for 30 seconds. The gels were developed in 3% sodium carbonate with 0.02% formaldehyde. The developed gels showing the pattern of protein bands was scanned and the image was analyzed.

### Western blotting

SDS/PAGE gels were transferred in Towbin buffer (25mM Tris, 192mM glycine, 20% w/v methanol) to polyvinylidene fluoride membrane, blocked in 1% bovine serum albumin (BSA) in PBS, incubated overnight at 4 degrees Celsius in a 1:1,000 dilution of 100ug/mL affinity-purified primary IGG to KAP-1, MTHFD1, hnrnpK, TUBB, or PDI in 1% BSA in PBS containing 0.1% Tween-20 (PBST). Membranes were then washed twice in PBST and incubated for two hours at room temperature in a 1:5000 dilution of secondary anti-rabbit or anti-mouse antibody coupled to alkaline phosphatase in PBST. Membranes were washed two more times in PBST then incubated in a developer solution prepared from 100 uL of 7.5 mg/mL 5-bromo-4-chloro-3-indolyl phosphate dissolved in 60% dimethyl formamide (DMF) in water and 100ul of 15 mg/ml nitro blue tetrazolium dissolved in 70% DMF in water, adjusted to 50mL with 0.1 Tris (pH 9.5) and 0.1 mM magnesium chloride. Membranes were scanned and the integrated density of protein band was measured on ImageJ. Averages and the standard deviation between repeated experiments were calculated and plotted on Microsoft Excel.

### Tandem mass spectrometry

Samples were processed by SDS PAGE using a 10% Bis-ttris NuPAGE gel with the MES buffer system. The mobility region was excised and washed with 25 mM ammonium bicarbonate followed by 15mM acetonitrile. Samples were reduced with 10 mM dithoithreitol and 60 degrees Celsius followed by alkylation with 5o mM iodacetamide at room temperature. Samples were then digested with trypsin (Promega) overnight (18 hours) at 37 °C then quenched with formic acid and desalted using an Empore SD plate. Half of each digested sample was analyzed by LC-MS/MS with a Waters NanoAcquity HPLC system interfaced to a ThermoFisher Q Exactive. Peptides were loaded on a trapping column and eluted over a 75uM analytical column at 350 nL/min packed with Luna C18 resin (Phenomenex). The mass spectrometer was operated in a data dependent mode, with the Oribtrap operating at 60,000 FWHM and 15,000 FWHM for MS and MS/MS respectively. The fifteen most abundant ions were selected for MS/MS.

Data was searched using a local copy of Mascot (Matrix Science) with the following parameters: Enzyme: Trypzin/P; Database: SwissProt Human (concatenated forward and reverse plus common contaminants); Fixed modification: Carbamidomethyl (C) Variable modifications: Oxidation (M), Acetyl (N-term), Pyro-Glu (N-term Q), Deamidation (N/Q) Mass values: Monoisotopic; Peptide Mass Tolerance: 10 ppm; Fragment Mass Tolerance: 0.02 Da; Max Missed Cleavages: 2. The data was analyzed by label free quantitation (LFQ) and spectral count methods. LFQ intensity values of each condition were measured in triplicate and compared against each other to generate log_2_ fold change values for each combination of conditions. Spectral counts were filtered for a 1% protein/peptide false discovery rate requiring 2 unique peptides per protein and the data set was further adjusted by subtraction of spectral counts for specific proteins observed in the control resin. Identified proteins were searched in the NURSA database of protein-protein interactions (https://dknet.org/about/NURSA_Archive) to determine if they interact with KAP-1, the Bushman labs oncogene database (http://www.bushmanlab.org/links/genelists) to determine if they were known to be implicated in cancer, and the VirusMentha database (https://virusmentha.uniroma2.it/) to determine if they interact with HIV. String diagrams of protein-protein interaction networks were generated using the STRING database (string-db.org).

## Acknowledgements

We thank the National Cancer Institute for running the sixty cell line screen described in Supplemental Figure 2. We thank Usha F. Lingappa for help with the figures, Dmitry Temnikov for IT support, Halley McCormick for help with data analysis, and Jairam R. Lingappa for valuable suggestions during manuscript preparation.

